# MRDadaptis: Self-Adaptive Parameter Configuration Enhances Minimal Residual Disease Detection in Heterogeneous ctDNA Samples

**DOI:** 10.1101/2025.08.20.671240

**Authors:** Tianci Wang, Xin Lai, Shenjie Wang, Zhengfa Xue, Yuqian Liu, Xiaoyan Zhu, Xiaonan Wang, Zhili Chang, Yang Shao, Xian Zhang, Jiayin Wang

## Abstract

Detection of structural variations (SV) through circulating tumor DNA (ctDNA) has become a key method for detecting minimal residual disease (MRD). However, the heterogeneity of ctDNA samples, characterized by variable limits of detection (LOD) and diverse structural variant types, significantly impacts detection stability and consistency, posing persistent challenges for conventional SV detection tools such as Delly and Manta. These widely-used methods require extensive manual parameter tuning, hindered by the combinatorial complexity of multiple parameters and heterogeneous sequencing data. To address this, we propose MRDadaptis, a novel SV detection tool that uniquely incorporates a self-adaptive parameter optimization mechanism. MRDadaptis distinguishes itself by integrating Bayesian optimization with meta-learning techniques to dynamically adjust detection parameters automatically, based on intrinsic features derived from the ctDNA sequencing data itself. This innovative approach not only reduces manual intervention but also effectively captures sample-specific characteristics, significantly improving detection stability and detection performance. Extensive validation experiments using both simulated and real-world ctDNA datasets demonstrates MRDadaptis’s distinct advantages, including markedly improved average F1-scores and superior stability (reduced variance, lower RMSE, increased kurtosis)

These results highlight the significant advantages of MRDadaptis in addressing sample heterogeneity, underscoring its potential to improve the accuracy and reliability of MRD detecting through ctDNA analysis.

https://github.com/aAT0047/MRDadaptis.git

## Introduction

Minimal residual disease (MRD) refers to the small number of cancer cells that remain in the body after treatment, which can provide valuable information for evaluating treatment effectiveness and assessing the risk of relapse [1]. Genetic variants, such as single nucleotide variants (SNVs) and insertions/deletions (indels), are identified in ctDNA and can serve as effective biomarkers for MRD detection [2–3]. Recently, structural variants (SVs) have gained increased recognition for their critical role in tumor initiation and progression, making SV detection a pivotal method for MRD assessment [4–5]. However, the performance of SV detection methods in ctDNA often demonstrates significant instability, posing a substantial challenge for accurate MRD detection [6–7].

MRD-positive status is generally defined by a variant allele frequency (VAF) threshold of 10□□, which requires a given mutation to be observed multiple times [8,9]. On the TEC-Seq platform, for example, hotspot mutations (VAF > 0.1%) must appear in at least three unique sequencing reads to count as MRD-positive, whereas non-hotspot mutations demand support from more than seven reads [10]. Thus, it is imperative for SV detection tools to maintain stable performance even in ctDNA with extremely low sequencing coverage. Unfortunately, existing SV detection tools frequently exhibit considerable performance fluctuations across heterogeneous tumor samples [11–14], leading to errors in MRD classification, such as high rates of false negatives (FN), causing missed MRD-positive diagnoses or false positives with potentially severe medical consequences. Although Studies have shown that optimizing parameters based on sample characteristics can significantly improve the performance of these tools [15]. the optimization of parameters for SV detection in ctDNA regions remains a complex challenge.

Why? Performance instability primarily arises from mismatches between sample-specific characteristics (e.g., insert size, base quality) and detection software parameters. Specifically, substantial heterogeneity exists among ctDNA samples regarding the limit of detection (LOD) for SVs and the variety of SV types. For instance, ctDNA levels in squamous cell carcinoma are generally higher compared to adenocarcinoma, causing significant variability in SV detection sensitivity across tumor types when employing uniform detection methods [16]. Moreover, SVs exhibit notable diversity in number, size (ranging from hundreds to millions of base pairs), type, and genomic location (including coding, non-coding, and regulatory regions), further complicating detection [17].

Therefore, achieving stable and precise MRD detection necessitates segmenting heterogeneous ctDNA regions and optimizing parameters specifically tailored to each region. However, these objective faces two significant challeng**es: (**□.) Parameter configurations often involve more than 10 highly coupled threshold parameters, making it exceedingly challenging to identify optimal configurations within a complex, high-dimensional parameter space. (□.) The vast number of cancer patients, extensive ctDNA base pairs, and inherent high heterogeneity introduce another significant obstacle. Each ctDNA sample requires segmentation into potentially thousands of subregions for independent parameter optimization, yet currently, parameter adjustments primarily rely on subjective, expert-driven manual tuning based on previous detection outcomes. Such manual tuning methods are highly subjective, inefficient, and lack reproducibility, especially when processing large numbers of heterogeneous samples [18–20].

To address these challenges, we propose MRDadaptis, a novel SV detection tool. MRDadaptis uniquely integrates split-read, read-pair, and read-depth strategies [21–23], automatically optimizing parameter configurations based on the inherent characteristics of heterogeneous sequencing regions. Unlike existing tools, MRDadaptis employs an innovative self-adaptive Bayesian framework [24], enabling automatic segmentation and parameter tuning tailored to specific ctDNA subregions. To enhance efficiency, MRDadaptis incorporates a meta-learning strategy [25–27] that leverages historical data for rapid, adaptive parameter recommendations in new ctDNA datasets.

MRDadaptis thus achieves precise quantification of sample characteristics and rapid optimization of detection parameters, providing stable, efficient, and accurate SV detection.

## Methods

MRDadaptis **(Fig. 1B)** enhances the standard SV detection workflow **(Fig. 1A)** by incorporating three additional steps: Data Segmentation, Bayesian optimization framework, and Meta Model. **As shown in Figure 1B**, the **data segmentation** module partitions the DNA sequencing data into multiple subregions, the **Bayesian framework** then maximizes the posterior probability of a surrogate model (Gaussian process) to determine the optimal parameters that minimize the cost function. It collects meta-features from sub-regions and parameter configurations (obtained via the Bayesian framework) as meta-targets, and uses these historical data to train the meta-model. Next, Next, the meta-model learns the mapping from features to meta-targets. This approach significantly reduces computational complexity and enables efficient and stable parameter recommendations. The core components of MRDadaptis are detailed below. **The specific algorithm is in the Supplementary Material. 2** (**Algorithm 2: MRDadaptis)**

**Figure 1:**
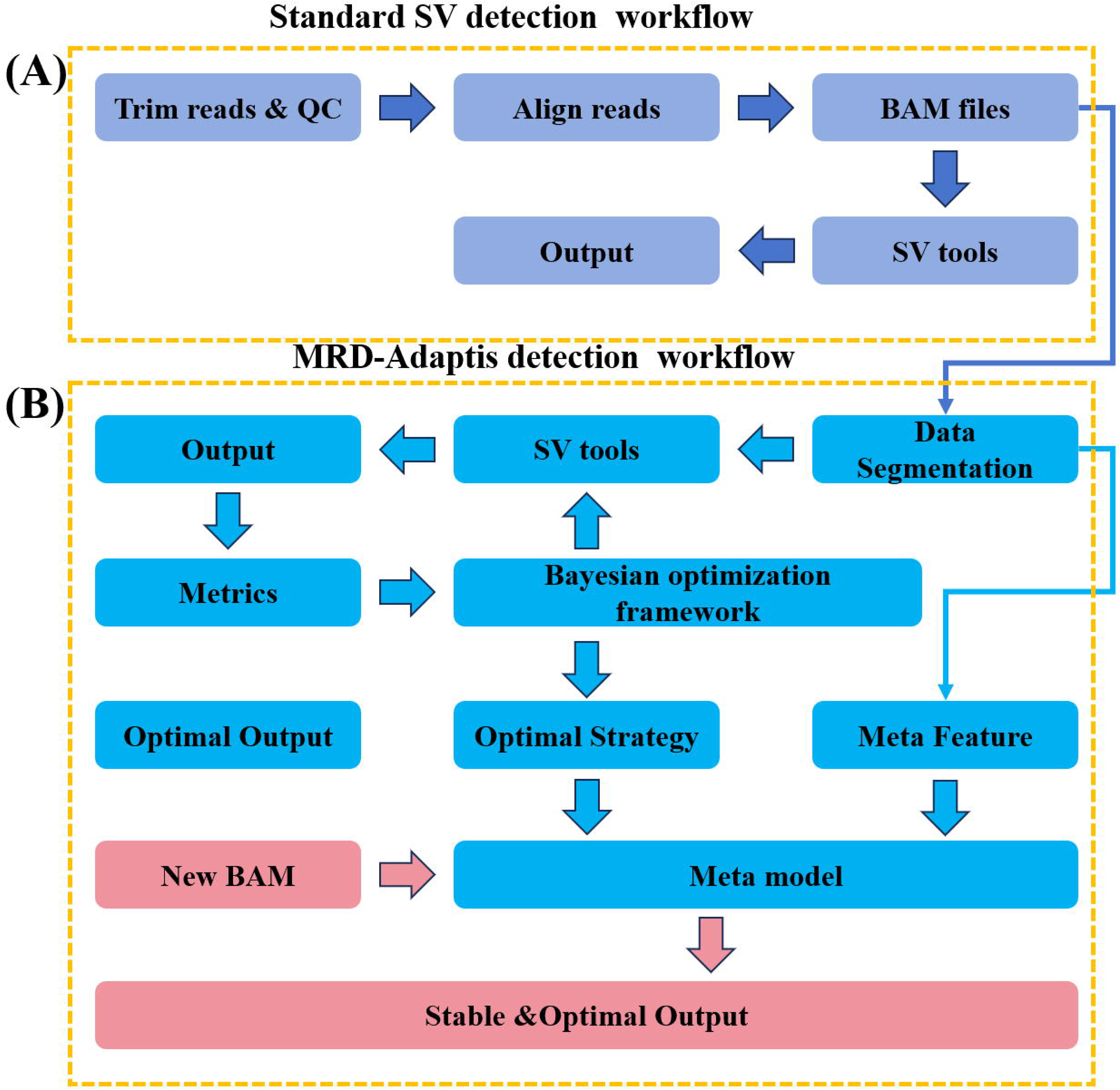
(**A**) Standard SV detection workflow. (**B**) Enhanced SV detection workflow with MRDadaptis, incorporating three additional steps: Data Segmentation, Bayesian Optimization Framework, and Meta Model, enabling adaptive parameter adjustments based on sample-specific characteristics to ensure stable SV detection results.

### Data Segmentation &Meta-feature extraction for sub-regions

During the data segmentation phase **(as shown in Figure 2)**, MRDadaptis employs an adaptive sliding window technique to scan the ctDNA data and identify structural variations (SVs). The window size is dynamically determined based on soft-clipping and mismatch information.

**Figure 2:**
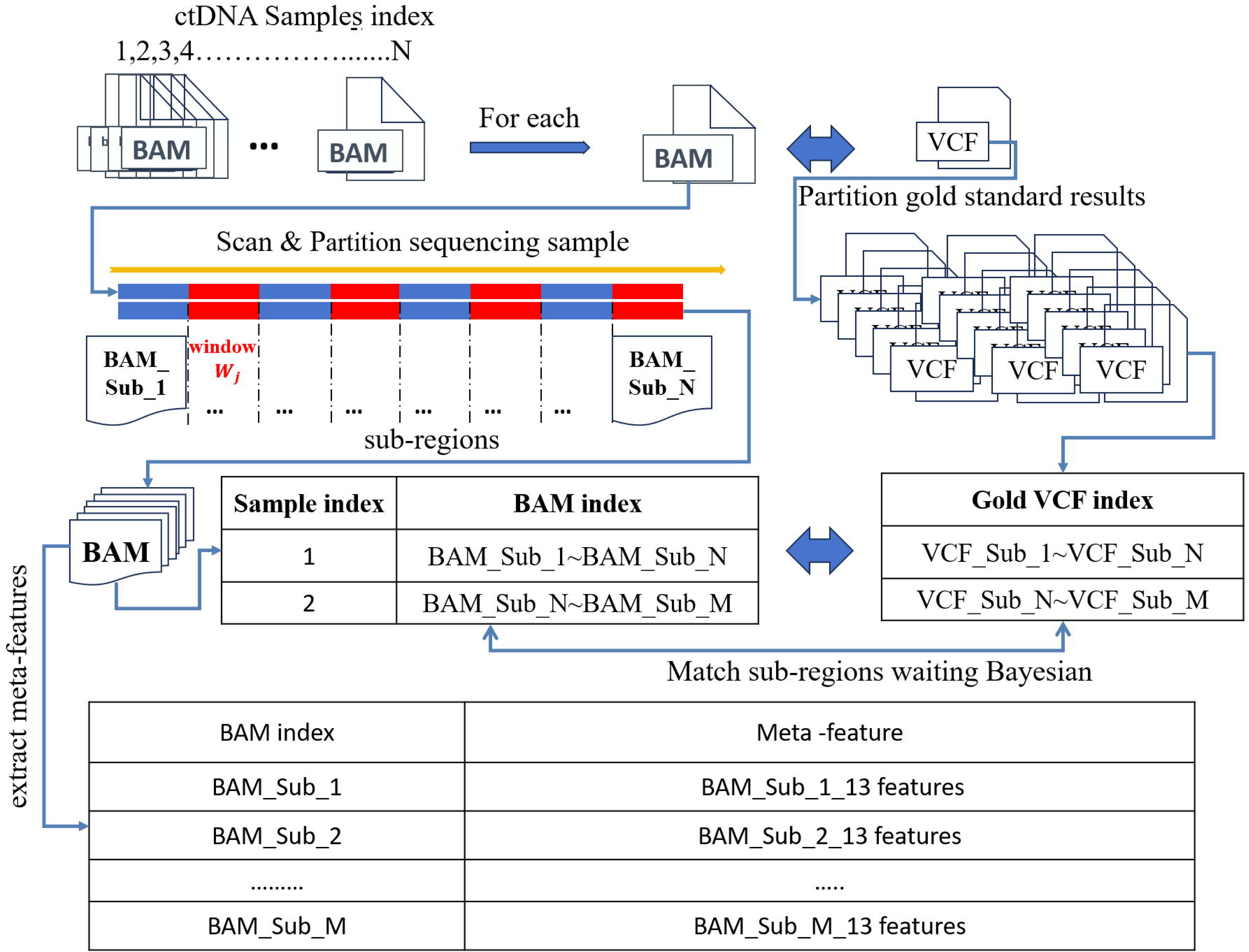
Data Segmentation and Meta-feature Extraction for ctDNA Sequencing Sub-regions. During the data segmentation phase, MRDadaptis utilizes a sliding window technique to scan the ctDNA sequencing sample, dividing it into sub-regions. After segmentation, the sub-regions are systematically renamed and renumbered to create new BAM file subsets, preserving their mapping relationship with the original sample. Meta-features are then extracted from the ctDNA data using tools like SAMtools and BCFtools. These meta-features, such as sequencing depth and mapping quality

### Step I: Adaptive Window Length Calculation

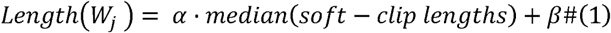

Where:

*W_j_* is the window size;

*α* and *β* are empirical parameters.

### Step II: Variation Score and Dynamic Threshold *T*

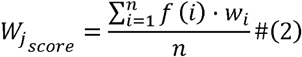

Where:

*f*(*i*) represents the variation value at position, (such as mismatch rate, etc.);

*w_i_* is the weight assigned to position, based on its relative distance from the center of the window;

*n* is the number of positions in the current window.

The dynamic threshold is *T* calculated based on the distribution of variation scores across the genome

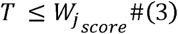

### Step □: Significant Variation Region Selection

A sub-region begin, end is considered to contain significant gene features if the following conditions are met:

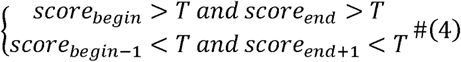

Finally, MRDadaptis extracts these sub-regions for further analysis. Throughout the splitting process, meticulous care is taken to prevent breakpoints from occurring in mismatched regions, ensuring the consistency and accuracy of each sub-region. Subsequently, these sub-regions are systematically renamed and renumbered to create new BAM file subsets (e.g., BAM_Sub_1, BAM_Sub_2, …, BAM_Sub_N) and establish mapping relationships with the original sample indices. Additionally, the gold standard results in the VCF files of these ctDNA data are segmented according to the sub-regions and correspond one-to-one with the respective sub-regions.

After obtaining the sub-regions containing variants, we extract meta-features that distinguish the calling performance of parameter configuration within these sub-regions. Inspired by the Wang et al. [20,38], we achieve this by extracting meta-features from the ctDNA data in BAM/SAM format files. Specifically, within the data segmentation module, we have integrated a fast-scanning algorithm, which differs from precise variant detection tools. This algorithm efficiently extracts meta-features by leveraging tools such as samtools, bcftools, and information from CIGAR, enabling rapid and effective feature extraction.

**As shown in Table 1**, this study, based on experimental results and a literature review [7–21], selects 13 meta-features to represent sub-regions of structural variations (SVs). For example, sequencing depth and mapping quality play a critical role in parameter optimization. Higher sequencing depth provides more accurate variant information, while higher mapping quality helps reduce false positive results. For low-depth or low-quality samples, we adjust the parameters to enhance the detection of shallow variants and mitigate the impact of alignment errors.

The combination of these features allows for effective differentiation of variants within ctDNA genomic regions, enabling the optimization of SV detection parameters for each sub-region. This approach minimizes redundancy and noise. The meta-features selected in this study have been specifically chosen due to their demonstrated effectiveness in distinguishing the performance of different parameter strategies across various genomic regions.

### Bayesian framework

To address the challenge of tuning over ten user-defined parameters required for SV detection on sub-regions (BAM file), where manual adjustment is infeasible due to the vast search space. MRDadaptis integrates a Bayesian optimization framework as the central driver for self-adaptive parameter optimization**. As shown in Figure 3**, the Bayesian Optimizer orchestrates the entire optimization cycle through a structured five-step process:

**Figure 3:**
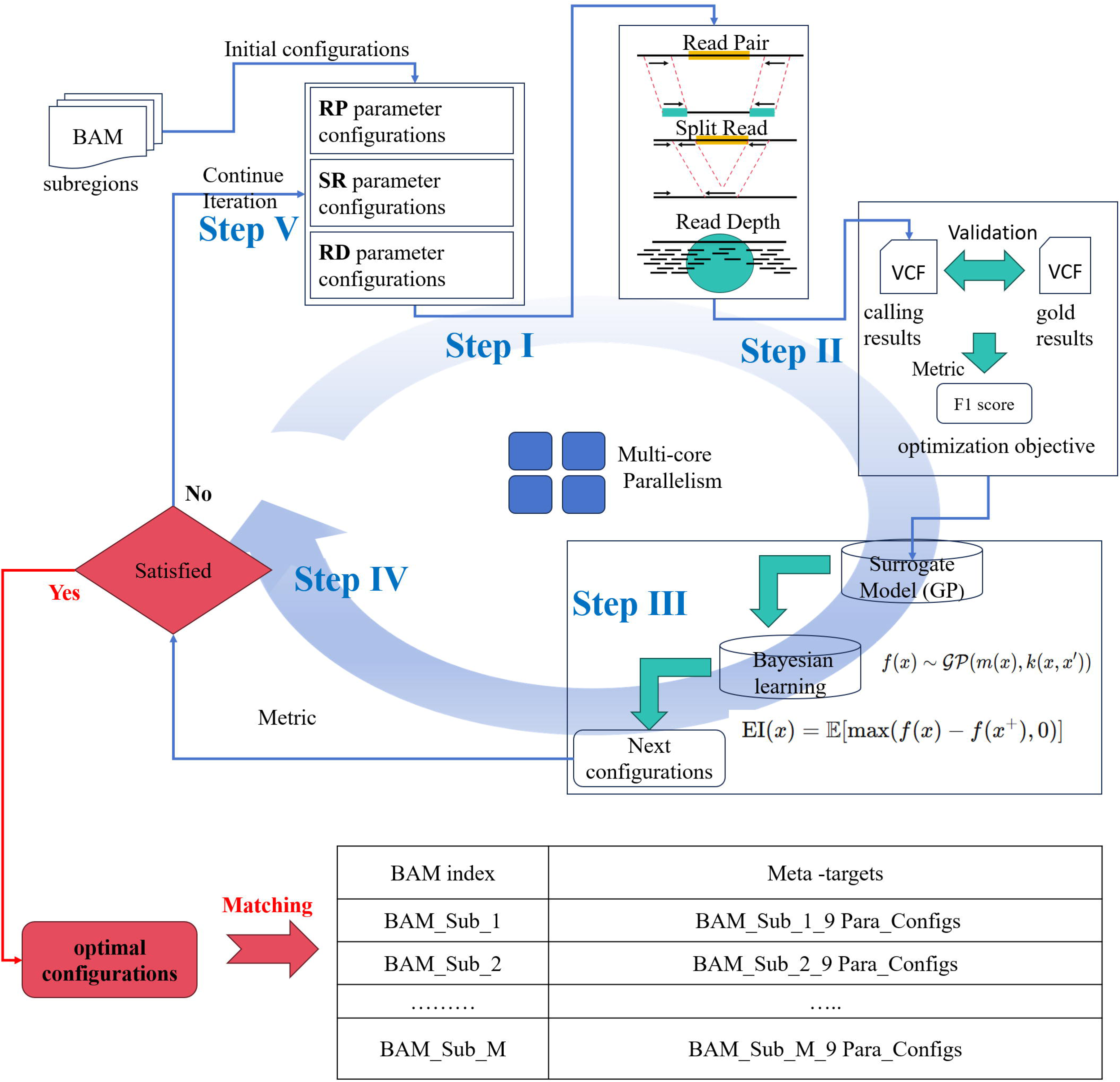
Bayesian Optimization Framework for Self-Adaptive Parameter Optimization in SV Detection: Step I – Initial SV Detection: Perform SV detection with initial parameters using read pair, read depth, and split read strategies. Step II – Evaluation: Evaluate VCF files against gold standard SV results and calculate performance metrics (e.g., F1 score). Step III – Surrogate Model: Construct a surrogate model using Gaussian Process (GP) for predictive modeling. Step IV – Acquisition Function: Maximize an acquisition function (e.g., Expected Improvement) to propose new parameter configurations. Step V – Iterative Updates: Refine the search space using Bayes’ Rule, gradually converging to optimal parameter settings.

**Step I:** Initial SV detection is performed using baseline parameter configurations, leveraging read pair, read depth, and split read strategies within a multi-core parallelism framework to generate preliminary SV results (VCF files).

**Step II:** The generated VCF files are evaluated against gold standard SV results to compute performance metrics (F1 score), which serve as feedback signals for Bayesian model updates.

**Step** □: Based on multiple sampled parameters sets and their corresponding detection performance metrics, a surrogate model *p*(*f*|*D_n_*) is constructed, where *D_n_* denotes the set of obsenved parameter-performance pairs. In the case of Gaussian process (GP) modeling the predictive posterior for any new configuration *x* is a normal distribution.

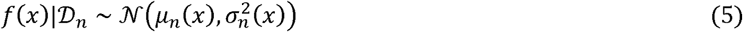

Where:

*f*(*x*):The (unknown) detection performance metric as a function of configuration *x*;

*D_n_*:The set of all observed pairs 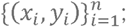

*μ_n_*(*x*):The posterior mean (expected value) of *f*(*x*) conditioned on the observations *D_n_*;

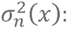 The posterior variance (uncertainty) of *f*(*x*) conditioned on the observations *D_n_*.

**Step IV:** Guided by the surrogate model, the Bayesian Optimizer proposes new parameter configurations by maximizing an acquisition function *ϕ*(*x*), such as Expected Improvement (EI)

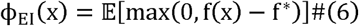

Where:

*ϕ*_EI_(*x*):Expected Improvement at configuration;

*f*(*x*):As above, the (unknown) detection performance at *x*;

*f* *:The best observed detection performance value so far;

E[•]: Expectation with respect to the predictive distribution of *f*(*x*).

**Step V:** Through iterative updates, the Bayesian Optimizer dynamically improves its internal model according to Bayes’ Rule:

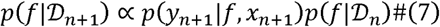

Where:

*p*(*f* | *D_n_*_+1_):Posterior distribution of *f* after observing *D_n_*_+1_.

*D_n_*_+1_ = *D_n_* ∪ {(*x_n_*_+1,_ *y_n_*_+1_)}

*p*(*y_n_*_+1_ | *f*, *x_n_*_+1_): Likelihood of observing *y_n_*+1 given *f* and *x_n_*+1.

*p*(*f* | *D_n_*):Prior (current) posterior distribution of *f* before the new observation.

progressively refining the search space and ultimately converging to an optimal set of parameter configurations for each sub-region. The optimization process terminates when either the performance metric (F1 score) exceeds 0.9, or the number of iterations reaches 500, whichever occurs first. This strategy effectively minimizes performance fluctuations across samples and ensures stable SV-based MRD detection

### The specific algorithm is in the Supplementary Material. 2 (Algorithm 1: Bayesian Optimization for Maximizing F1 Score)

Traditional SV evaluation metrics often define true positives (TP) as having more than 0.8 overlap with the reference standard, but this approach inherently introduces false positives. In the context of Bayesian optimization, such traditional evaluation criteria may fail to clearly distinguish between the global optimum and suboptimal solutions, especially in the early exploration phases of the objective function. When the function’s variation is minimal or unclear, the model may overlook subtle fluctuations, causing it to converge prematurely into local optima. This phenomenon, known as the flat region problem, arises when the objective function shows slow or minimal changes in certain regions, making it difficult for the optimization algorithm to reliably identify the global optimum. To mitigate this issue, we have implemented stricter evaluation standards designed to increase the model’s sensitivity to slight variations in the objective function, thus reducing the likelihood of the optimization process getting stuck in local optima prematurely.

The TP formula is as follows:

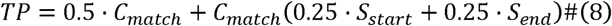

Where:

*C_match_* : chromosome matching.

*C_match_* = 1, If the chromosomes match,

*C_match_* = 0, otherwise.

This part contributes 50% (0.5) of the total TP score.

*S_start_* : Similarity score for the predicted and reference starting positions, calculated as:

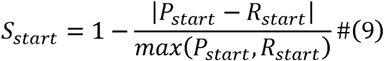

*S_end_* : Similarity score for the predicted and reference ending positions, calculated as:

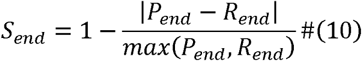

Where:

*P_start_ :* Predicted start position of the structural variant.

*P_end_ :* Predicted end position of the structural variant.

*R_start_:* Reference (true) start position of the structural variant.

*R_end_*: Reference (true) end position of the structural variant.

### Meta-Model

In our previous work, during the Data Segmentation phase, we partitioned the ctDNA subregions. Subsequently, in the Bayesian framework phase, we matched optimal parameter configurations to these heterogeneous window regions. These sets of parameter configurations served as labels or meta-targets in the meta-learning approach **(Table 2)**. Combined with the corresponding regional meta-features **(Table 1)**, 10-fold cross-validation was employed to divide the data into training, testing, and validation sets, which were used to train and evaluate the meta-model.

As illustrated in **Figure 4**, we employed this meta-model, which maps dataset features to parameter strategies with the objective of minimizing the historical loss function *L*(*f*) .

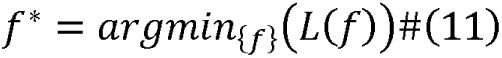

**Figure 4:**
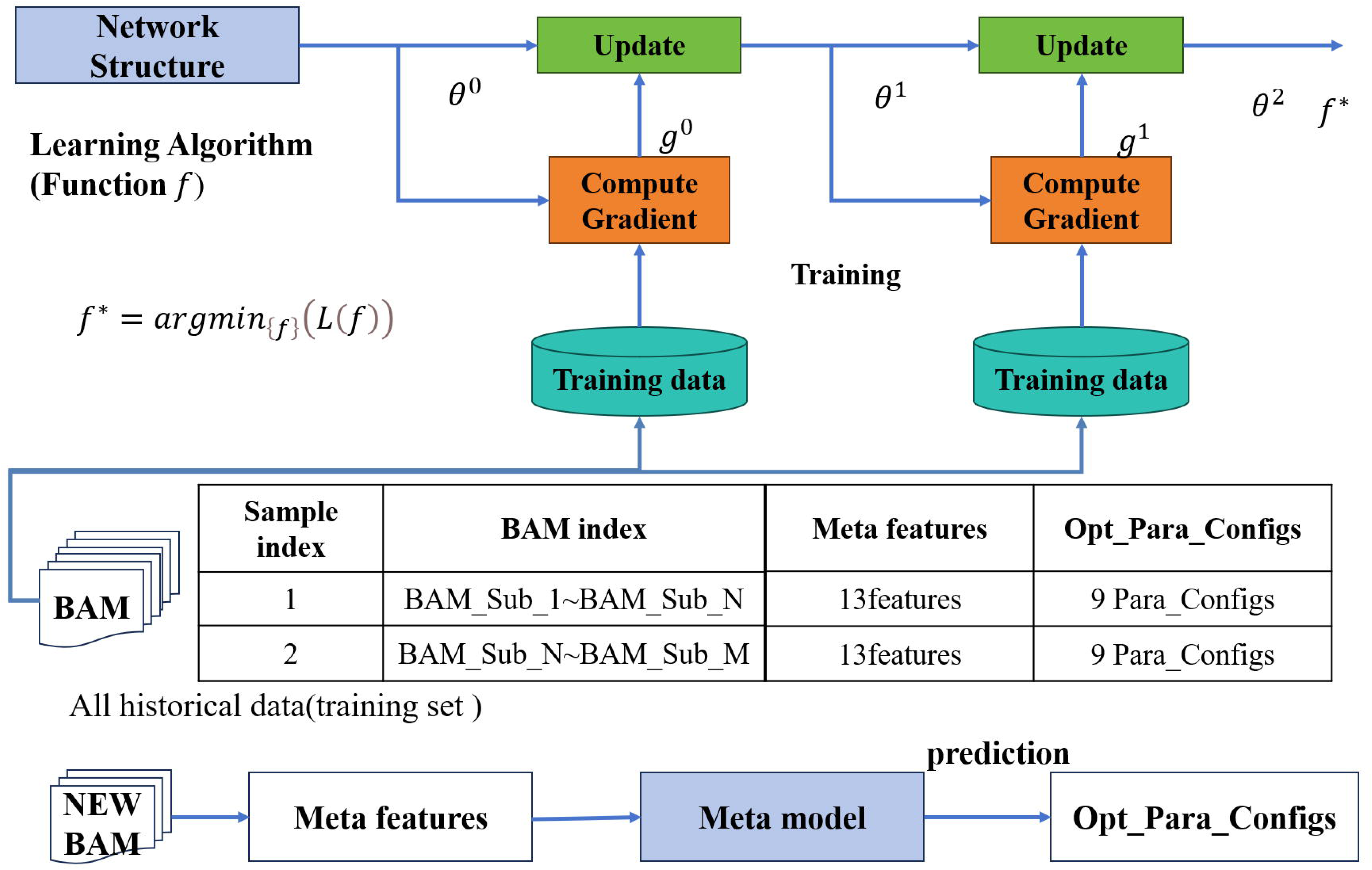
The meta-model leverages trained heuristic mappings to predict optimal parameters. In the data segmentation phase, meta-features for each subregion are extracted (Table 1). In the Bayesian phase, the optimal parameter configurations for each subregion are generated as meta-targets (Table 2). These meta-targets, combined with the corresponding meta-features, are used to train the meta-model. The meta-model then predicts the optimal parameter configuration for new subregions to optimize SV detection performance. The approach is validated using 10-fold cross-validation to ensure robustness and generalization to unseen data.

Where:

*f*:A candidate parameter strategy mapping function;

*f* *:The optimal parameter strategy function that minimizes the loss;

*L*(*f*):The historical loss function associated with mapping function *f*.

For the parameter recommendation problem, we define a meta-learning function as follows:

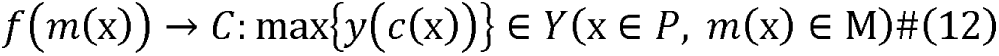

Where:

*P*: The set of all sequencing data requiring parameter recommendation;

*x*: An individual sequencing dataset;

*m*(*x*):The meta-features extracted from data *x*, *m*(*x*) ∈ *M*;

*M*:The set of all possible meta-feature vectors;

*f*(*m*(*x*)): The meta-model that maps meta-features to parameter strategies *c*:The space of all candidate parameter strategies;

*c*:The chosen parameter strategy, *c* ∈ *C*;

*y*(*c*(*x*)): The performance function that evaluates strategy *c* on data *x*;

*Y*:The space of possible performance values.

When new ctDNA sequencing data require parameter recommendations, users first extract meta-features from the regions (SAM/BAM files). These features are then input into the model, which outputs parameter recommendations aligned with specified meta-targets.

## Results

In the results section, we conducted an extensive evaluation of eight variant callers—Delly [28], Lumpy [29], Manta [30], BreakDancer [31], Pindel [32], MetaSV [33], SvABA [34], and MRDadaptis. We analyzed the stability of SV detection results across heterogeneous samples. We performed rigorous testing on both simulated and real datasets, which demonstrated the superior detection performance and stability of MRDadaptis in variant detection, particularly in the complex genomic scenarios of ctDNA research. We ensured that the validation datasets were distinct from the training datasets to guarantee unbiased and robust results. **For detailed descriptions refer to the final section of the supplementary materials.4.**

### Materials

This study utilized a total of 600 simulated ctDNA samples, 38 real-world ctDNA samples from IGGC/ POG-CA 176 real-world ctDNA samples from Nanjing Gene Sequencing Technology Co., Ltd. The simulated samples were generated using GSDcreator software [35], These heterogeneous ctDNA samples were generated by varying sequencing depth, variant type, variant length, and tumor concentration.Specifically,during the simulation process, structural variants (SVs) including deletions, inversions, translocations, and duplications were randomly introduced at intervals ranging from 1,000 bp to 1,000,000 bp, with SV sizes meticulously set between 100 bp and 25,000 bp. To emulate the characteristics of ctDNA, such as low tumor purity and DNA fragmentation, sequencing reads from the “tumor” and “normal” genomes were mixed in varying proportions. Specifically, we established multiple concentration layers at 0.01%, 0.05%, 0.1%, 0.2%, 0.5%, and 1% to encompass the range of ctDNA concentrations commonly observed in clinical samples.Regarding sequencing parameters, each sample was sequenced to a depth between 2000x and 5000x to address the low abundance of ctDNA, ensuring sufficient coverage for the detection of low-frequency variants. Paired-end reads were generated with lengths ranging from 100 bp to 150 bp to better simulate the highly fragmented nature of ctDNA.

The first set of real ctDNA samples were provided by Nanjing Gene Sequencing Technology Co., Ltd. and were collected between January 2023 and June 2024. Selected samples included Soft Tissue Sarcoma (STS), Lung Cancer (LC), and Lymphoma (Lymph) cases, with tumor content diluted to 0. 1-1% VAF to reflect the typical abundance of ctDNA in plasma. High-depth gene sequencing was performed using the Shi He No.1 panel to ensure sufficient coverage for detecting low-frequency variants in ctDNA. The sequencing included samples with detected gene fusions to capture structural variations relevant to ctDNA analysis. A total of 53 STS samples, 70 LC samples, and 53 Lymph samples were included in the analysis. These real samples were sequenced in FASTQ format and aligned using the Burrows-Wheeler Aligner (BWA) to ensure accurate mapping to the reference genome file hs37d5, also known as b37 + decoy, released by the 1000 Genomes Project (Phase II). All participants provided written informed consent prior to participation, ensuring the ethical conduct of the study and safeguarding the informed rights of the participants.

The second set of real ctDNA samples were provided by the BC Cancer’s Personalized OncoGenomics (POG) program and are accessible through the International Cancer Genome Consortium’s Accelerating Research in Genomic Oncology (ICGC-Argo) platform (URL: https://platform.icgc-argo.org/)Normal samples were subsampled to the GRCh38DH reference genome (GRCh38 with decoy and HLA sequences). Reads from tumor and matched normal samples were computationally mixed using a depth-adjusted subsampling approach. SAMtools randomized read selection function (-s parameter) was used to generate variant allele frequencies (VAFs) in the range of 0.01%–0.1%. Access to raw sequencing data and clinical metadata is restricted to authorized researchers through the ICGC-Argo platform, in compliance with ethical guidelines and donor consent agreements. Data requests require institutional review board (IRB) approval and project-specific authorization from the ICGC-Argo Data Access Committee.

The third set of real ctDNA samples were provided by the Pancreatic Cancer Harmonized “Omics” Analysis for Personalized Treatment (PACA-CA) program, accessible via the ICGC-Argo platform (URL: https://platform.icgc-argo.org/).To establish a high-confidence structural variation (SV) detection panel, we utilized detection data from 152 pancreatic ductal adenocarcinoma (PDAC) cases from the PACA-CA initiative. The workflow was as follows:

All structural variant calls were first aggregated and then filtered through four layers of orthogonal criteria:(i) Retention only of variants passing BRASS internal quality filters;(ii) Minimum structural variant length of 50 bp;(iii) Breakpoint homology ≤ 2 bp;(iv) Localization exclusively to primary reference chromosomes (chr1–22, X). After stringent filtering, we obtained a final high-confidence SV panel consisting of 2,771 unique breakpoint connections **(see Supplementary Table S7)** [36]. Subsequently, the resulting multi-sample VCF file was converted into BEDPE format.

**The MRD testing cohort** was generated as follows: (i) We selected CRAM files from 20 tumor samples in the PACA-CA dataset; (ii) Matched these with 20 normal samples randomly selected from the Utah Residents with Northern and Western European Ancestry (CEU, 1000 Genomes Project **see Supplementary Table 6**); (iii)Used computational methods to merge tumor and normal samples at variant allele fractions (VAF) of 0.01%-0.1%, emulating clinically relevant low-level circulating tumor DNA (ctDNA) signals to serve as MRD-positive samples; (iv)The same 20 normal samples from the 1000 Genomes Project were utilized individually as negative controls, ensuring the absence of tumor signals.

### Metrics to evaluate the detection performance and stability

In each ctDNA dataset, the performance of MRDadaptis was initially evaluated using precision, recall, and F1-score for individual samples. However, assessing detection performance alone is insufficient due to inherent heterogeneity across genomic regions within samples. To ensure reliable MRD detection, it is critical to maintain stable performance across these heterogeneous genomic regions. Therefore, additional statistical metrics—including mean, standard deviation (std), coefficient of variation (CV), root mean square error (RMSE), kurtosis, and skewness— were introduced specifically to quantify the stability of MRDadaptis results. Consequently, our evaluation strategy clearly distinguishes between two categories of metrics: those evaluating detection performance (precision, recall, F1-score), and those evaluating the stability and consistency of this performance across heterogeneous genomic regions (mean, std, CV, RMSE, kurtosis, skewness). **The specific equation is in the Supplementary Material 3 (Metrics to evaluate the performance).**

### SV detection on simulated data

We evaluated the **performance and stability** of MRDadaptis and other SV calling tools in SV detection using simulated data. To assess performance (Table 3), we analyzed recall, precision, and F1 scores,which were calculated based on the mean values from the test sample set. To evaluate stability, we examined the standard deviation (Std) of F1 scores and the coefficient of variation (CV) of F1 scores. These metrics allowed us to distinguish between the performance and stability of each tool’s performance across different sample types.

**For performance,** MRDadaptis performed exceptionally well(F1 score= 0.89), far surpassing other tools, especially Pindel (F1 score =0.44). This indicates that MRDadaptis maintains high performance in variant detection and can sensitively detect SV signals even under low VAF conditions in ctDNA.

**For stability,** we measured performance using the standard deviation (Std) and coefficient of variation (CV) of F1 scores. Delly (recall = 0.63), Lumpy (recall = 0.64), and Pindel (recall = 0.75) demonstrated higher sensitivity in certain samples but exhibited significant performance fluctuations in more complex samples. For instance, Pindel (F1_CV =0.64), indicating that the performance of the caller is closely related to sample characteristics. When detection parameters do not align with sample characteristics, performance inevitably fluctuates.

In contrast, based on the adaptive parameter optimization mechanism, MRDadaptis (F1_CV=0.23) indicates stable performance across heterogeneous samples. Even in highly heterogeneous regions of ctDNA scenarios, MRDadaptis maintains consistent detection performance, which is beneficial for the stability of MRD detection.

### SV detection on real-world data

We further evaluated MRDadaptis using real data, demonstrating exceptional stability and superior performance. In a dataset of 176 samples from Nanjing Gene Sequencing Technology Co., Ltd., we used MRDadaptis’ Bayesian framework module to output the optimal parameter configuration based on the gold-standard results for LC **(Supplemental Table 2)** and Lymph datasets**(Supplemental Table 3**). The features extracted from these two datasets, combined with the parameter configurations, were used as historical data to train the meta-model**(Supplemental Table 7**). The STS dataset was then used as a test set to validate the performance of MRDadaptis.

We compared the metrics(Mean,Std &CV) of multiple structural variant detection tools on STS data—including DELLY, LUMPY, Manta, BreakDancer, Pindel, MetaSV, SvABA, and MRDadaptis**(Table4)**. MRDadaptis consistently maintained **high performance**(F1=0.99,precison =1,recall=0.98) and **stable**(e.g.F1,mean=0.99,std=0.03) Metrics, highlighting its strong ability to handle sample heterogeneity(STS,**supplemental_table1**).

To further demonstrate the generalization ability of the MRDadaptis meta-model module, we selected 38 samples from the POG-CA project as the second test set. We evaluated the impact of SV types on the performance of different callers under the condition of VAF = 0.1% **(Table 5)**. Compared to other callers, MRDadaptis showed stable performance across all SV types and exhibited high F1 scores (F1_mean = 0.89).To further highlight the stability of MRDadaptis, we evaluated the performance of 8 callers on samples with VAF values of 0.01%, 0.02%, 0.04%, 0.06%, 0.08%, and 0.10% **(Table 6)**. The concentration of VAF was positively correlated with the detection performance of most callers. However, MRDadaptis maintained stable detection performance across multiple VAF concentrations. At VAF = 0.01%, the detection performance of all callers sharply declined, but MRDadaptis still performed the best (F1_score = 0.37).

These results demonstrate that the MRDadaptis meta-model exhibits good generalization capability. Even in unrelated cohorts, the meta-model can quickly configure parameters based on sample characteristics, ensuring stable SV detection and laying a foundation for MRD detection.

### Potential usage in MRD detection

In evaluations of 20 MRD-negative and 20 MRD-positive samples, MRDadaptis achieved the lowest combined error rate(**Fig.5A**) and maintained high precision (≥0.92) at recall levels ≥0.9 in precision–recall analyses(**Fig.5B**). Crucially, according to the UpSet plots (**Fig.5**C/D), only four SV breakpoints were unanimously reported by all eight callers in the MRD-negative cohort, whereas in MRD-positive samples MRDadaptis consistently detected ≥4 high-confidence breakpoints in every sample, while other methods typically identified only two to three. These findings demonstrate MRDadaptis’s superior sensitivity and robustness for ultra-low-frequency (10^-4^ LOD) SV detection.

**Figure 5:**
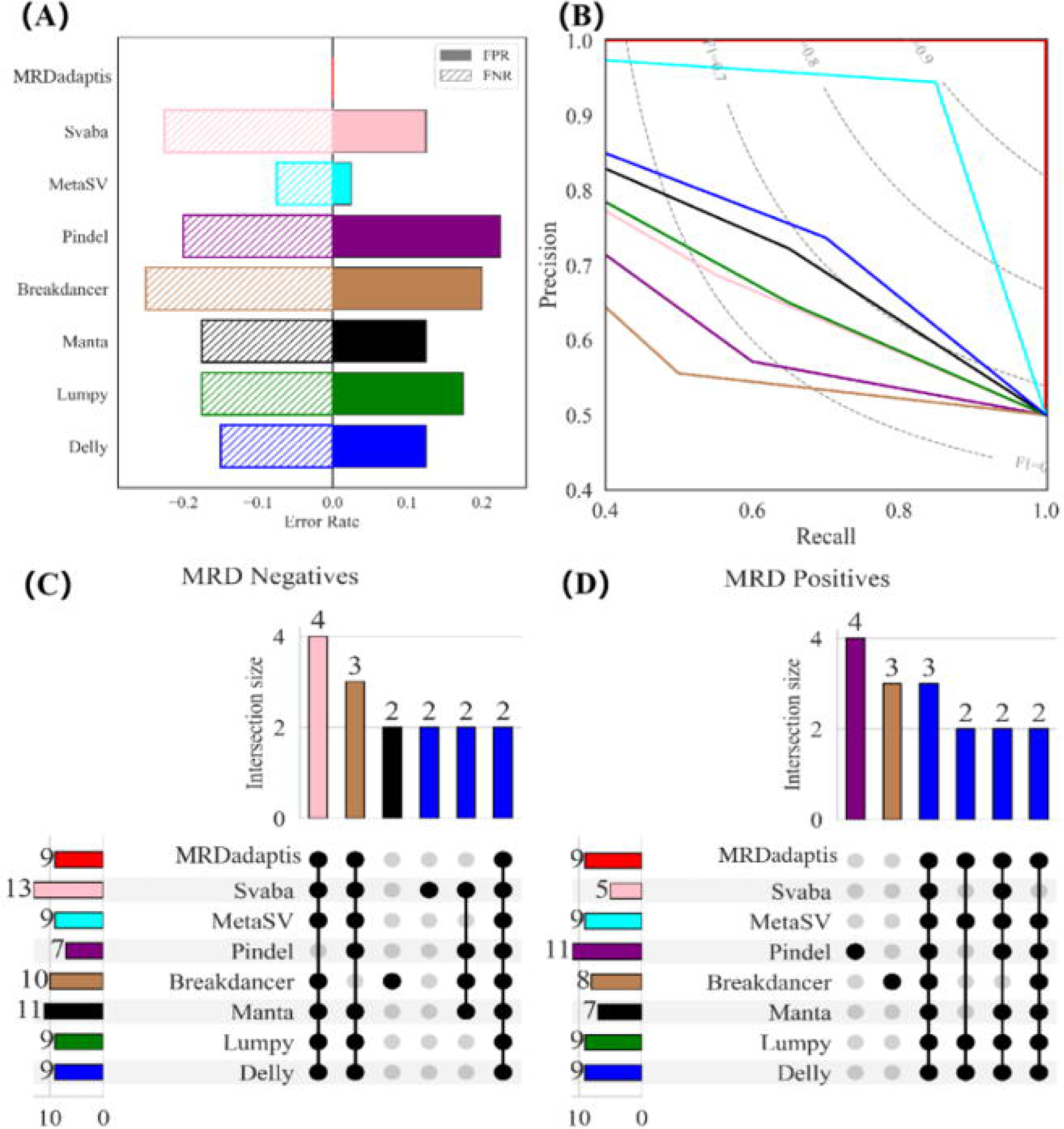
Performance of MRDadaptis versus seven existing SV callers in MRD detection.**(A)** Total error rate (Error Rate = FPR + FNR): grey bars denote false positive rate (FPR), hatched bars denote false negative rate (FNR).**(B)** Precision–Recall curves; dashed grey lines indicate F1-score contours.**(C)** Upset plot for MRD-negative samples: shows intersection sizes of breakpoint calls among eight tools; the top bars indicate the number of tools in each intersection.**(D)** Upset plot for MRD-positive samples at 10^-4^ LOD: MRDadaptis detects ≥ 4 panel SVs in every positive sample, outperforming all other callers.

### Why MRDadaptis Can Achieve Stable SV Detection in Heterogeneous ctDNA Samples?

To further demonstrate that targeted parameter configurations can effectively stabilize SV detection results, we performed a detailed analysis of the meta-features from 173 real datasets provided by ShiHe Genomics. The Bayesian framework was employed to optimize parameters based on the maximum F1 score as the objective. Parameter optimization was performed for each sub-region, and the iterative process was terminated when the F1 score exceeded 0.9 or after 500 iterations. These optimal configurations were used as meta-targets and combined with meta-features from the Data Segmentation module to form a historical training dataset for the meta-model. MRDadaptis was able to adaptively optimize parameters for each sample region.

Using the Lymph sample set as an example **(additional data available in the supplementary materials.4 e.g. LC,STS.)**, **as shown in Fig.6A**, we selected the top five features based on standard deviation for analysis: Read Mapping Bias (RMB), Average Sequencing Depth, Insert Size Mean, Insert Size Standard Deviation (Insert Size Stdev), and High Mean Density Percentage(HMDP). These features were normalized to allow for comparison on the same scale (**supplemental Table4**). Significant variation exists among the features across samples, with some features exhibiting extreme values exceeding 0.9. For example, **as shown in Fig.6B**, tools like LUMPY and DELLY exhibited performance instability in samples with large insert size variability. Variability in Read Mapping Bias (RMB) and sequencing depth also impacted the performance of other tools like BreakDancer and Pindel. However, MRDadaptis demonstrated high adaptability by adjusting its detection parameters based on sample characteristics, maintaining stable performance despite feature fluctuations.

**Figure 6:**
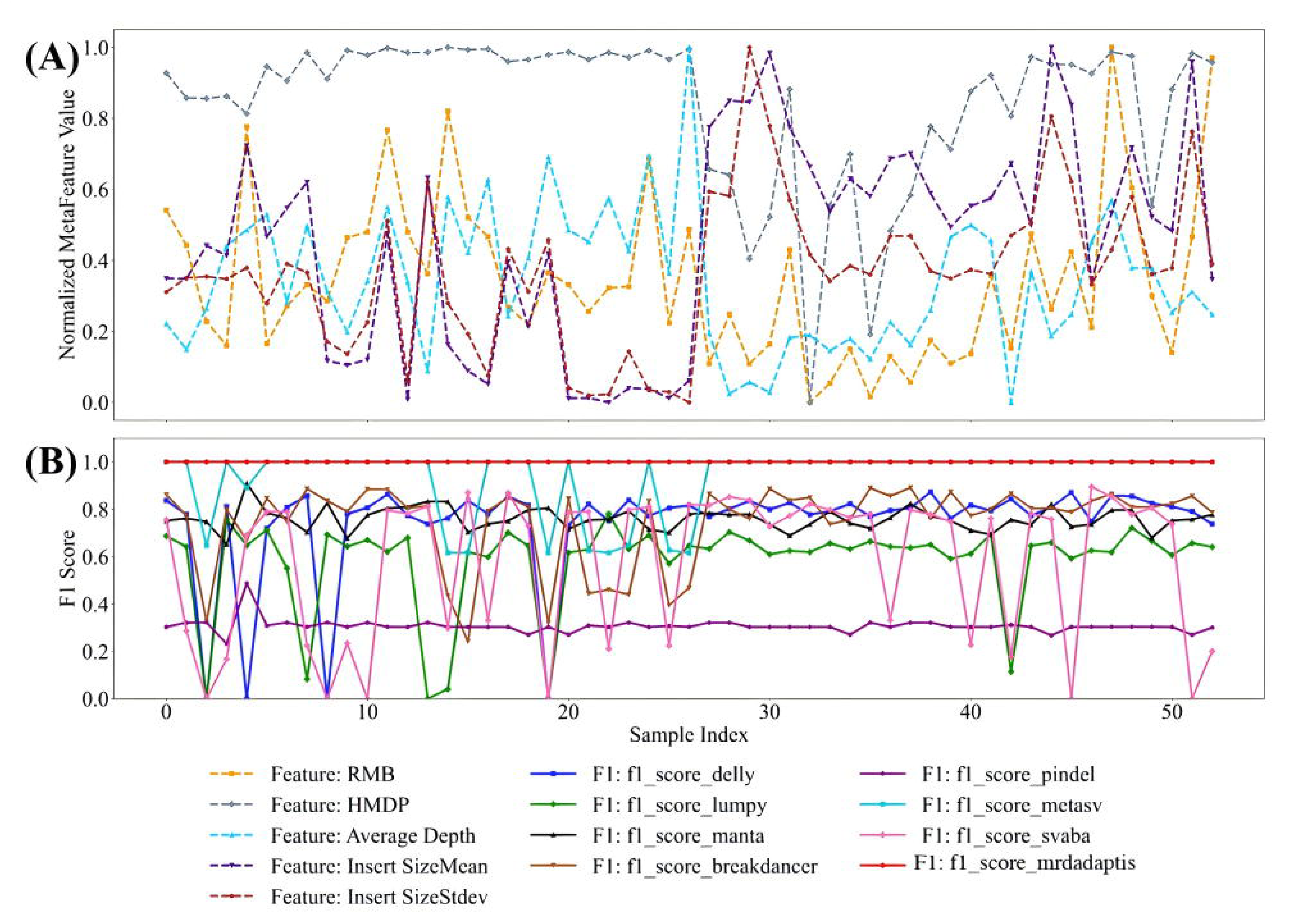
Lymph Sample Analysis(A) This panel highlights the significant heterogeneity in lymph sample features, focusing on the five features with the highest variability: mapping bias, sequencing depth, insert size, insert size standard deviation, and mutation density. These features show substantial fluctuations across samples and are normalized to a common scale for comparison. (B) Feature fluctuations impact the performance and stability of SV detection tools. Tools such as LUMPY and DELLY, which rely on insert size or sequencing depth, exhibit noticeable performance variation under changing feature conditions. High mapping bias introduces noise, reducing the effectiveness of tools like BreakDancer and Pindel. Low sequencing depth and elevated mutation density further complicate detection, particularly for tools optimized for high-confidence calls. Despite these challenges, MRDadaptis maintains high detection performance and stability across diverse feature conditions.

Different tumor datasets exhibit higher heterogeneity, resulting in greater fluctuations in sample features. In further analysis with 176 samples from three datasets (LC, Lymph, and STS) **.As shown in Figure 7**,MRDadaptis exhibited a unimodal F1-score distribution, tightly concentrated between 0.9 and 1.0, reflecting consistency and stability across diverse samples. In contrast, other tools like Delly and Manta showed more dispersed F1-score distributions, with some samples failing to detect variants. Tools like Lumpy and BreakDancer showed lower F1 scores, highlighting poor consistency. Pindel’s F1-score distribution was multimodal, reflecting significant fluctuations across samples.

**Figure 7:**
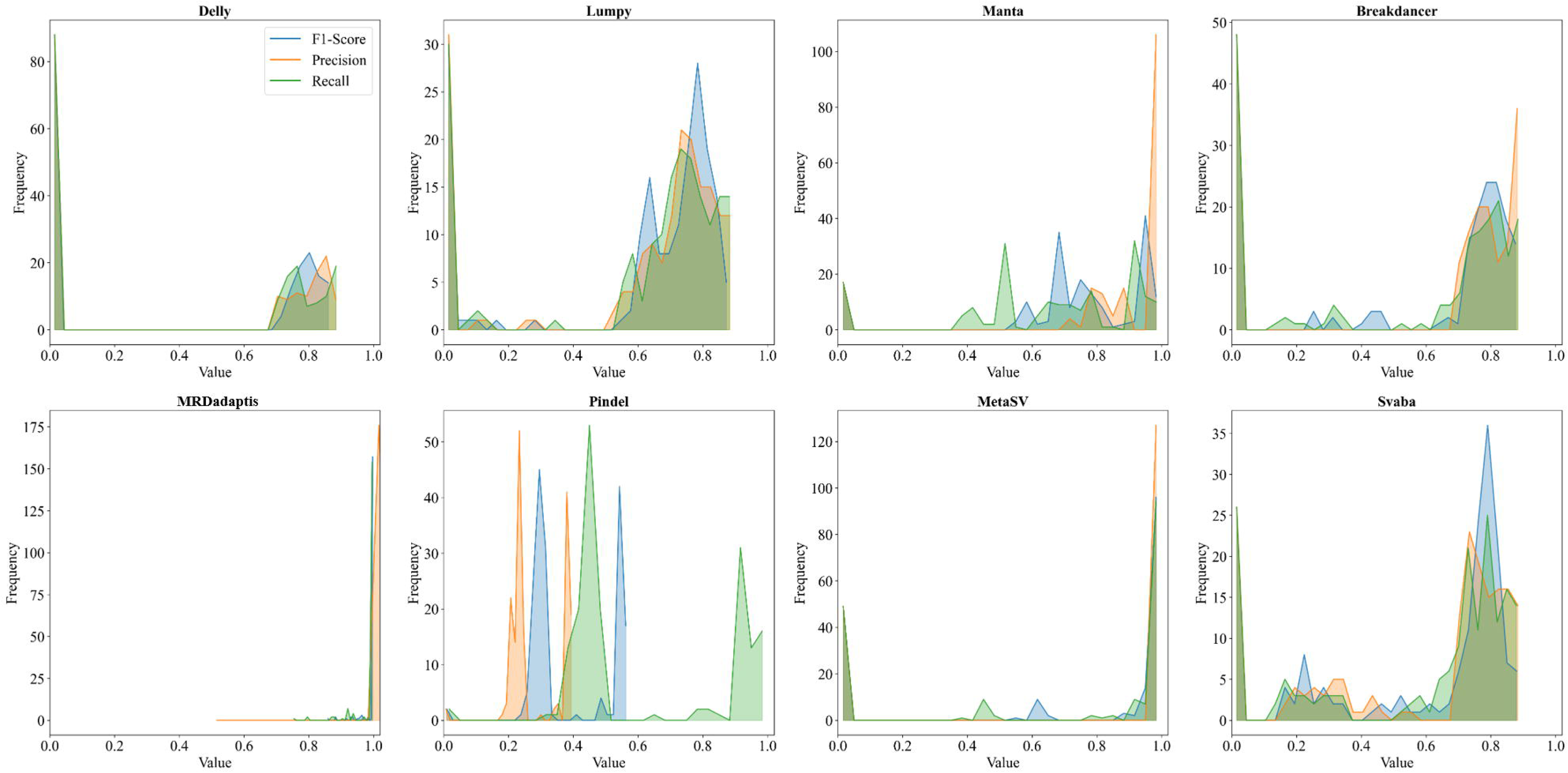
F1-Score Distribution Across Different Tumor Datasets.This figure shows the F1-score distribution of various SV detection tools across three tumor datasets (LC, Lymph, and STS) comprising 176 samples. MRDadaptis displays a unimodal and tightly concentrated distribution, indicating consistently high performance and stability across diverse samples. In contrast, tools like Delly and Manta present more dispersed distributions, with some samples failing to detect variants. Lumpy and BreakDancer exhibit generally lower F1 scores, reflecting poor consistency. Pindel shows a multimodal distribution, suggesting substantial performance fluctuations across the dataset.

Furthermore, **as depicted in Figure 8**, We performed a joint analysis of Root Mean Square Error (RMSE), Skewness, and Kurtosis for the F1-scores of the tools. MRDadaptis achieved an exceptionally low RMSE (0.02196), indicating that it consistently achieved optimal performance. Additionally, the negative skewness (-4.137) and high kurtosis (18.2717) of MRDadaptis’s F1-score distribution showed that its performance was concentrated in the high-performance region, with minimal outliers **(supplemental Table5)**.

**Figure 8:**
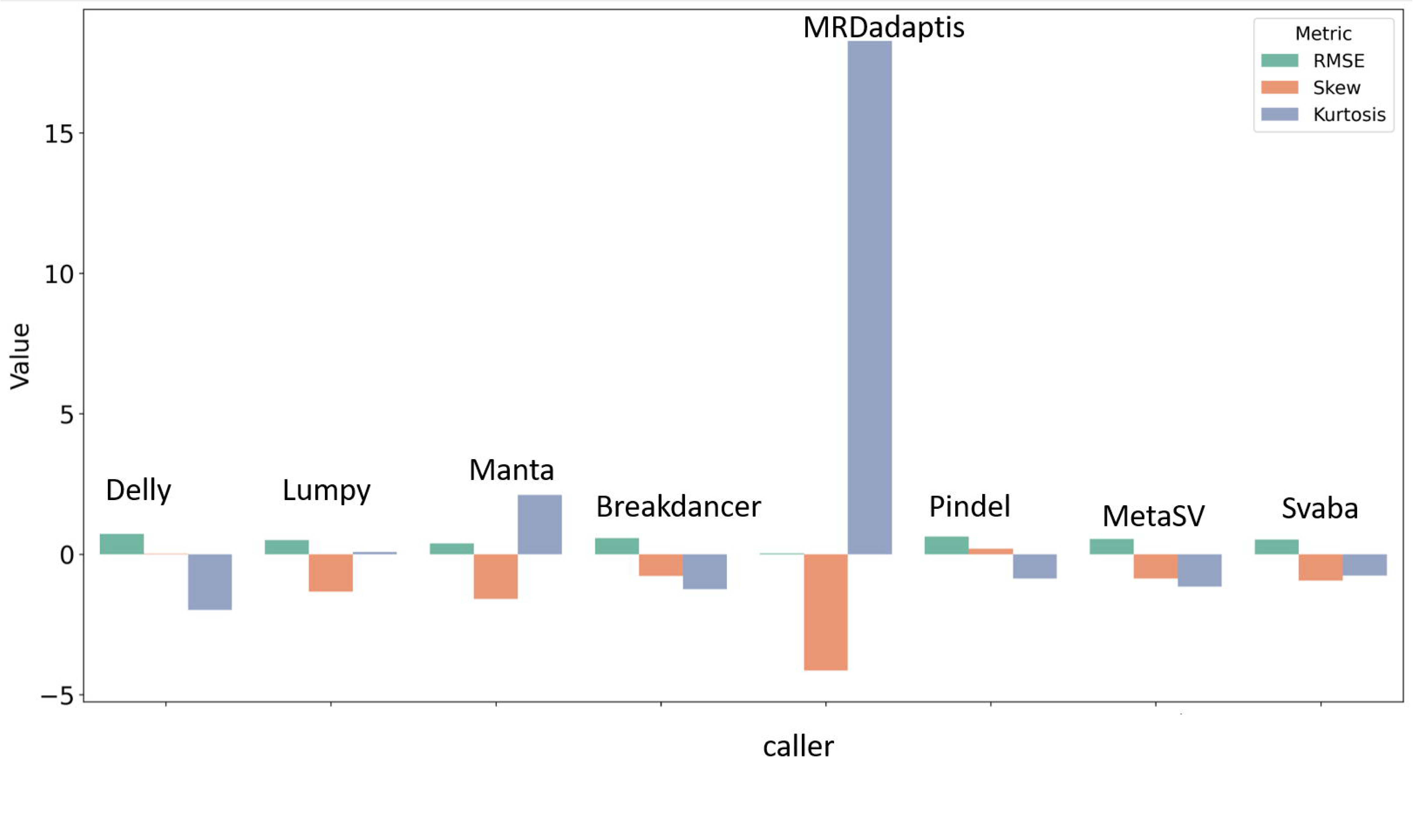
Comparative analysis of SV detection tools based on the F1-scores of 176 real-world samples. Metrics include the RMSE of F1-scores relative to the optimal performance (F1=1) and the skewness and kurtosis of normalized F1-score distributions.

In contrast, other tools exhibited more varied distributions and higher skewness and kurtosis values, reflecting their instability in handling heterogeneous samples. Pindel and MetaSV, for example, showed multimodal distributions and uneven score distributions, highlighting their struggles in maintaining consistent performance across different sample characteristics.

Specifically,tumor samples have higher SV density and more complex types; fixed parameter settings can lead to increased false positives and false negatives. This is because the complexity of different samples requires the parameters in detection strategies to be adaptively adjusted to accurately capture true variant signals.However, parameters related to detection strategies such as split read, read pair, and read depth need adaptive adjustment across different samples. For example, the mean and standard deviation in the read pair strategy need to be adjusted according to the insert size distribution of the sample; the search range parameters in the split read strategy need to adapt to different genomic complexities; the thresholds in the read depth strategy need to consider the sample’s coverage and noise levels. Fixed parameter settings can lead to suboptimal detection performance, potentially missing true variants or introducing false positives due to sequencing or alignment errors. Therefore, achieving adaptive parameter adjustment is crucial for SV detection in different samples [37–38].

### Why Mitigating Bayesian Optimization Framework Bottlenecks with Meta-Modeling is Important?

Bayesian optimization frameworks can iteratively optimize parameters based on gold-standard results, yielding the optimal configuration. However, there are two key bottlenecks:□Bayesian optimization process requires multiple rounds of initial data collection and iteration to achieve optimization, resulting in high time complexity.□Bayesian optimization requires a defined optimization target,but with new sequencing data, the optimization target is unknown, impeding iterative optimization.To address these issues, we propose a meta-learning module. By leveraging historical data with gold-standard results, the Bayesian module is first used to determine the optimal parameter configuration as the meta-target. The meta-model can quickly recommend parameter configurations for new data based on its features, enabling generalized and adaptive parameter optimization.

We randomly selected 100 samples from real-worldand simulated datasets (100 testing dataset) for testing. The results demonstrate that the meta-model leverages the meta-dataset to rapidly identify optimal parameter configurations, efficiently minimizing the loss function without unnecessary exploration. **As illustrated in Figure 9**, the meta-model reduces the average parameter recommendation time to 25 seconds, significantly outperforming Bayesian optimization. This acceleration is attributed to the meta-model’s ability to swiftly predict optimal settings, thereby minimizing redundant exploration. Furthermore, **Table 7** shows that the meta-model maintains high performance and stability in variant detection while enhancing computational efficiency. These findings highlight the effectiveness of integrating meta-learning with Bayesian optimization for complex genomic data analysis.

**Figure 9:**
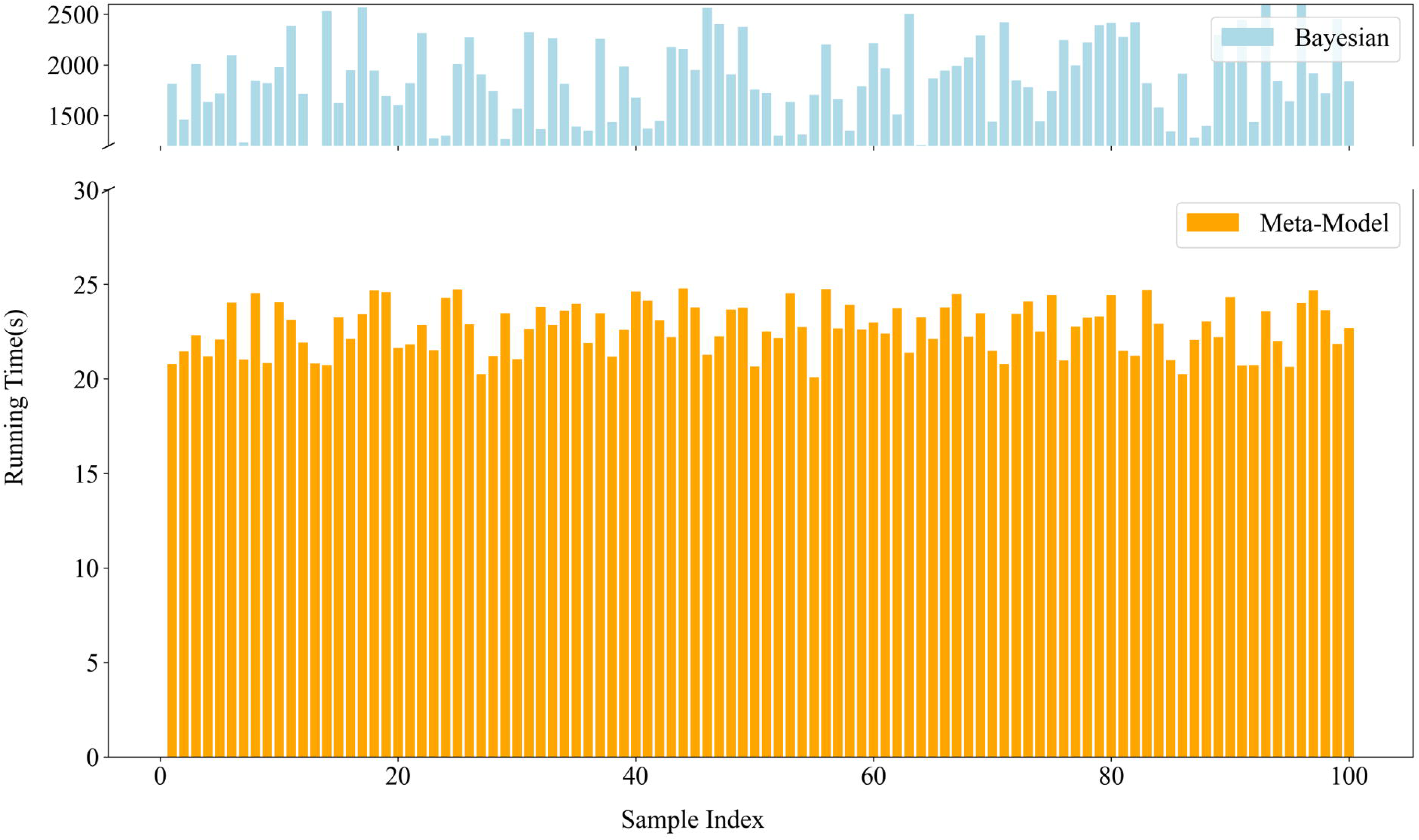
Comparison of the runtime(s) between the meta-model and Bayesian optimization frameworks for parameter tuning, highlighting the significant efficiency improvement achieved by the meta-model.

## Discussion

In this study, we first demonstrated that the parameter-tuning problem for structural variant (SV) callers can be rigorously formulated as a Boolean satisfiability (SAT) problem, proving its NP-hardness **(supplementary materials.1)**. This underscores the impracticality of exhaustive search within high-dimensional parameter spaces, particularly in large and heterogeneous ctDNA cohorts. To overcome this computational bottleneck, we developed a Bayesian optimization framework leveraging Gaussian process uncertainty quantification, efficiently guiding the parameter search towards promising regions of the parameter space. Furthermore, MRDadaptis integrates meta-learning derived from historical optimization data, effectively “warm-starting” the Bayesian search. This significantly reduces computational demands associated with repeated SV calling, thereby enabling adaptive and sample-aware parameter recommendations unattainable through brute-force or heuristic methods alone **(supplementary materials.6).**

To further justify the choice of meta-features utilized in MRDadaptis, we systematically evaluated their contributions to SV-calling performance through SHAP (SHapley Additive exPlanations) value analysis **(Fig. 10)**. Specifically, sequencing quality indicators (e.g., Average Depth, Average Read Length) and SV complexity metrics (e.g., HMDP, Short SV Percentage, Repeat Percentage) exhibited consistently high SHAP values across model outputs. These findings highlight the necessity and effectiveness of these meta-features in capturing critical variations relevant to adaptive parameter tuning.

**Figure 10:**
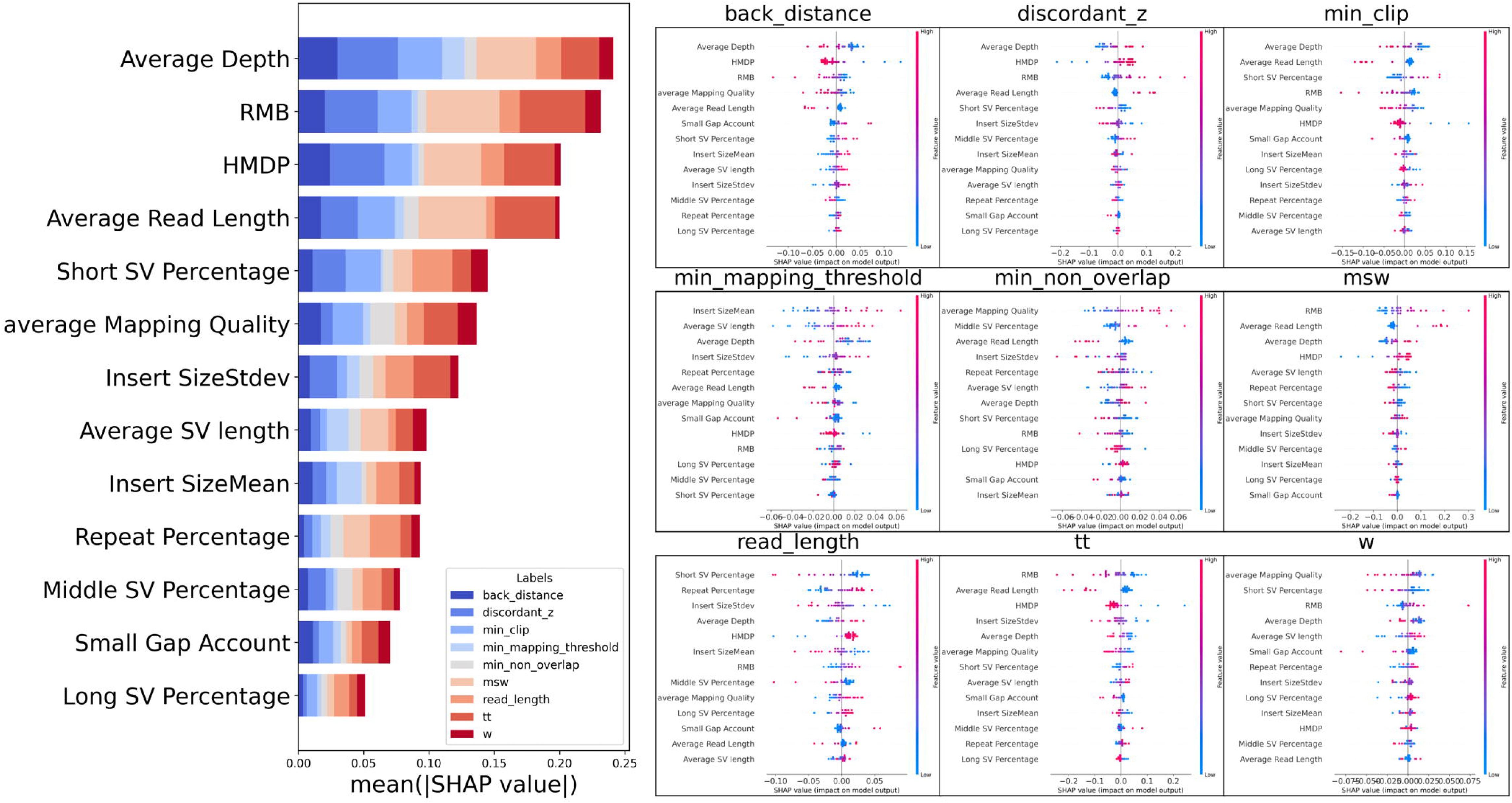
SHAP-based interpretability analysis of meta-feature importance across SV-calling parameters. The left panel ranks the 13 selected meta-features by their mean absolute SHAP values, reflecting their overall impact on parameter optimization. Sequencing quality features (e.g., Average Depth, Read Length) and SV complexity metrics (e.g., HMDP, Short SV Percentage) show the highest contributions. The right panel presents SHAP summary plots for individual SV-caller parameters, illustrating how each feature influences model output. Red points indicate higher feature values; blue points indicate lower ones.

Finally, recognizing that accuracy and stability are paramount for therapeutic decision-making and prognostic assessment in clinical minimal residual disease (MRD) monitoring, we conducted a comprehensive stability analysis **(Fig. 11)** [39–40]. Specifically, root mean square error (RMSE) directly reflects the quantitative accuracy and repeatability of MRD detection. Lower RMSE values indicate a robust ability to consistently and precisely detect minimal residual disease at extremely low variant allele frequencies (VAF), significantly enhancing sensitivity to subtle tumor burden fluctuations. Meanwhile, skewness and kurtosis provide complementary insights into measurement reliability: skewness identifies systematic biases toward consistent over- or underestimation, which could lead to overtreatment or missed diagnoses if significantly deviated from zero; kurtosis evaluates the risk of sporadic extreme errors that could compromise the stability and reliability of longitudinal clinical monitoring [41–42]. Thus, an ideal MRD detection platform must not only deliver high sensitivity and specificity but also demonstrate a favorable “low RMSE, near-zero skewness, and low kurtosis” profile. This comprehensive stability ensures accurate, consistent, and clinically actionable MRD quantification, enabling precise individualized treatment adjustments and reliable long-term patient management.

**Figure 11:**
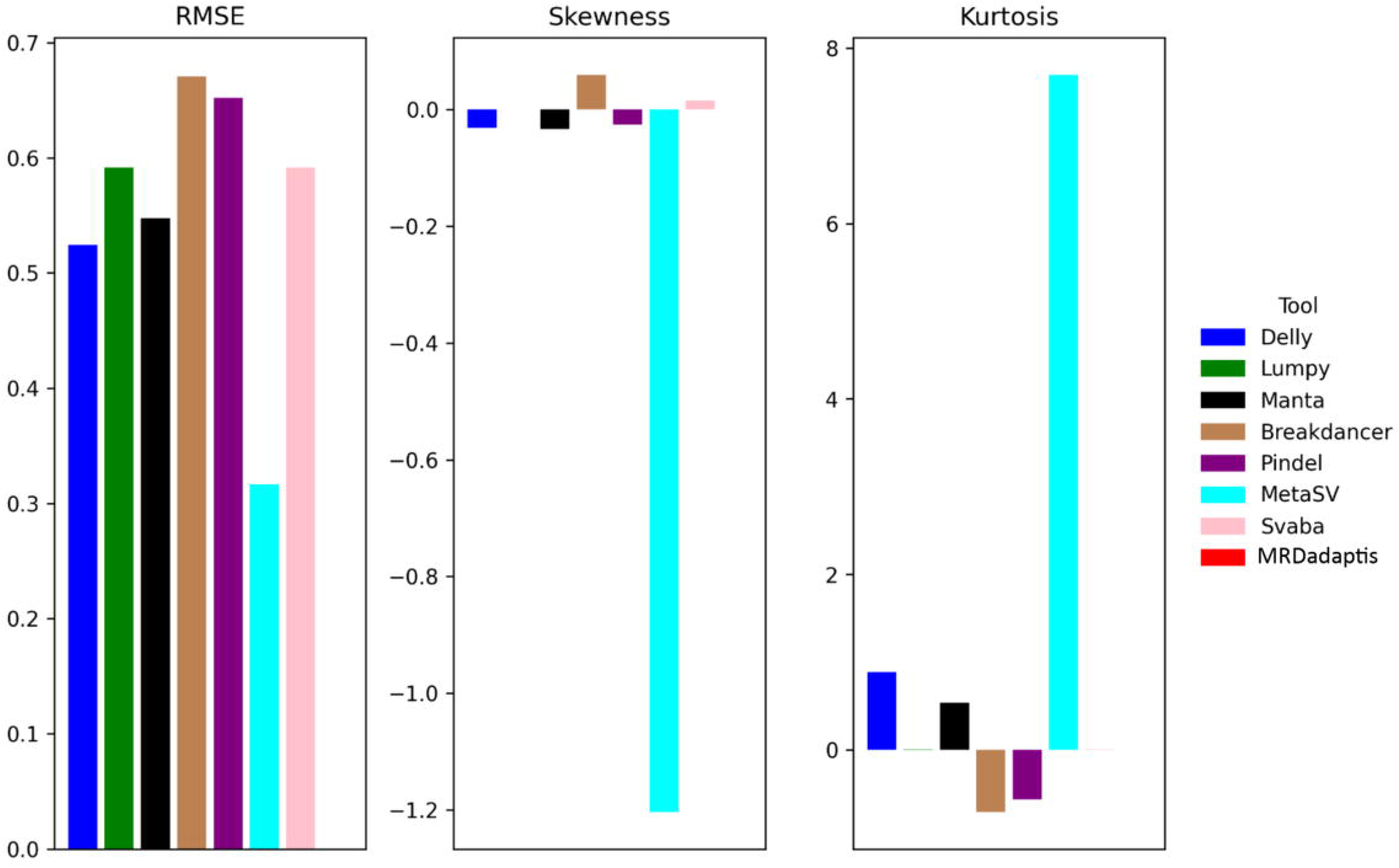
Stability analysis of SV detection tools at clinically defined MRD thresholds, as measured by root-mean-square error (RMSE), skewness, and kurtosis.

## Conclusion

MRDadaptis provides a robust, adaptive solution for SV detection in ctDNA, overcoming both the statistical heterogeneity of tumor samples and the computational inefficiency of traditional parameter search. The integration of Bayesian optimization with meta-learning ensures rapid convergence to near-optimal settings, yielding unimodal F1-score distributions tightly centered between 0.9 and 1.0. As such, MRDadaptis stands to improve the reliability of MRD monitoring in clinical settings and offers a generalizable framework for precision genomic analyses.

### Key Points

**MRDadaptis is introduced as a novel tool** designed to ensure high detection performance and stable structural variant (SV) detection performance across heterogeneous ctDNA regions, overcoming the limitations of traditional SV detection methods

**MRDadaptis incorporates a self-adaptive mechanism** that optimizes detection parameters based on the specific characteristics of ctDNA sequencing data, ensuring consistent performance across varied sample types

**MRDadaptis distinguishes itself by integrating Bayesian optimization and meta-learning** to dynamically adjust detection parameters for each sub-region in ctDNA data, providing a more flexible and adaptive approach compared to existing methods that rely on uniform parameter settings.

**MRDadaptis effectively mitigates SV detection performance fluctuations,** ensuring more accurate MRD detection.

**Extensive validation demonstrates that MRDadaptis achieves superior performance metrics**. The advantages of MRDadaptis are validated through extensive experiments on both simulated and real-world ctDNA datasets, demonstrating its superior stability, improved performance metrics, and enhanced reliability in MRD detection. .

## Supplementary data

Supplementary data is available at Briefings in Bioinformatics online.The algorithm can be accessed at https://github.com/aAT0047/MRDadaptis.git for academic usage only.

## Author contributions

J.W. and T.W. conceived and designed this research; J.W. and W.T. X.Z. designed the model; Z.X., T.W. and X.Z.implemented the program and performed the experiments; X.W., Y.S. and Z.Z. provided and analyzed the data. T.W., J.W. wrote the manuscript. Z.X., Y.L.,L.L.and T.W. conducted the revision. All authors have read and agreed to the latest version of the manuscript.

## References

1. Elsea D, Carlson JJ, Eckert B, et al. MRD Testing in Multiple Myeloma: Modeling the Potential Clinical and Economic Outcomes Based on the Master Trial.□Blood. 2024;144:5028.

2. Barris DM, Weiner SB, Dubin RA, et al. Detection of circulating tumor DNA in patients with osteosarcoma.□Oncotarget. 2018;9(16):12695.

3. Liu Y, Jiang T, Gao Y, et al. Psi-caller: a lightweight short read-based variant caller with high speed and accuracy[J]. Frontiers in Cell and Developmental Biology, 2021, 9: 731424

4. Su J, Yang X, Zhang L, et al. 506P Development of a sensitive urinary cfDNA profiling assay for detection and therapy efficacy monitoring in bladder cancer.□Ann Oncol. 2024;35:S1588–S1589.

5. Schütte J, Reusch J, Khandanpour C, et al. Structural variants as a basis for targeted therapies in hematological malignancies.□Front Oncol. 2019;9:475303.

6. Lee AY, Wong KF, Saffari A, et al. Combining accurate tumor genome simulation with crowdsourcing to benchmark somatic structural variant detection. Genome Biol. 2018; 19: 1–15.

7. Hu H, Gao R, Gao W, et al. SVDF: enhancing structural variation detect from long-read sequencing via automatic filtering strategies[J]. Briefings in Bioinformatics, 2024, 25(4): bbae336.

8. Nagafuji K, Miyamoto T, Eto T, et al. Monitoring of minimal residual disease (MRD) is useful to predict prognosis of adult patients with Ph-negative ALL: results of a prospective study (ALL MRD2002 Study)[J]. Journal of hematology & oncology, 2013, 6: 1–7.

9. Killock D. Early MRD predicts disease recurrence and benefit from adjuvant chemotherapy in CRC[J]. Nature Reviews Clinical Oncology, 2023, 20(3): 137–137.

10. Abbosh C, Birkbak N J, Swanton C. Early stage NSCLC—challenges to implementing ctDNA-based screening and MRD detection[J]. Nature Reviews Clinical Oncology, 2018, 15(9): 577–586.

11. Gong T, Hayes VM, Chan EKF. Detection of somatic structural variants from short-read next-generation sequencing data. Brief Bioinform. 2021; 22(3): bbaa056.

12. Gao R, Hu H, Jiang Z, et al. SVHunter: long-read-based structural variation detection through the transformer model[J]. Briefings in Bioinformatics, 2025, 26(3): bbaf203.

13. Zhang Z, Jiang T, Li G, et al. Kled: an ultra-fast and sensitive structural variant detection tool for long-read sequencing data[J]. Briefings in Bioinformatics, 2024, 25(2): bbae049.

14. Cameron DL, Di Stefano L, Papenfuss AT. Comprehensive evaluation and characterisation of short read general-purpose structural variant calling software. Nat Commun. 2019; 10(1): 3240.

15. Schikora-Tamarit M À, Gabaldón T. PerSVade: personalized structural variant detection in any species of interest[J]. Genome Biology, 2022, 23(1): 175.

16. Kosugi S, Momozawa Y, Liu X, et al. Comprehensive evaluation of structural variation detection algorithms for whole genome sequencing. Genome Biol. 2019; 20: 1–18.

17. Moding EJ, Nabet BY, Alizadeh AA, et al. Detecting liquid remnants of solid tumors: circulating tumor DNA minimal residual disease.□Cancer Discov. 2021;11(12):2968–2986. Brady L, Kriner M, Coleman I, et al. Inter- and intra-tumor heterogeneity of metastatic prostate cancer determined by digital spatial gene expression profiling. Nat Commun. 2021; 12(1): 1426.

18. Chen PS, Zeng ZY. Developing two heuristic algorithms with metaheuristic algorithms to improve solutions of optimization problems with soft and hard constraints: An application to nurse rostering problems. Appl Soft Comput. 2020; 93: 106336.

19. Bischl B, Binder M, Lang M, et al. Hyperparameter optimization: Foundations, algorithms, best practices, and open challenges. Wiley Interdiscip Rev Data Min Knowl Discov. 2023; 13(2): e1484.

20. Wang S, Liu Y, Wang J, et al. Is an SV caller compatible with sequencing data? An online recommendation tool to automatically recommend the optimal caller based on data features. Front Genet. 2023; 13: 1096797.

21. Tang K, Wang W, Sun Y, et al. Prophage Tracer: precisely tracing prophages in prokaryotic genomes using overlapping split-read alignment. Nucleic Acids Res. 2021; 49(22): e128–e128.

22. Tischler G, Leonard S. biobambam: tools for read pair collation based algorithms on BAM files. Source Code Biol Med. 2014; 9: 1–18.

23. Kadalayil L, Rafiq S, Rose-Zerilli MJJ, et al. Exome sequence read depth methods for identifying copy number changes. Brief Bioinform. 2015; 16(3): 380–392.

24. Wang X, Jin Y, Schmitt S, et al. Recent advances in Bayesian optimization. ACM Comput Surv. 2023; 55(13s): 1–36.

25. Zhang K, Chen N, Liu J, et al. An efficient meta-model-based method for uncertainty propagation problems involving non-parameterized probability-boxes. Reliab Eng Syst Saf. 2023; 238: 109477.

26. Schuler R. An algorithm for the satisfiability problem of formulas in conjunctive normal form. J Algorithms. 2005; 54(1): 40–44.

27. Lake BM, Baroni M. Human-like systematic generalization through a meta-learning neural network. Nature. 2023; 623(7985): 115–121.

28. Rausch T, Zichner T, Schlattl A, et al. DELLY: structural variant discovery by integrated paired-end and split-read analysis.□Bioinformatics. 2012;28(18):i333–i339.

29. Layer RM, Chiang C, Quinlan AR, et al. LUMPY: a probabilistic framework for structural variant discovery.□Genome Biol. 2014;15:1–19.

30. Chen X, Schulz-Trieglaff O, Shaw R, et al. Manta: rapid detection of structural variants and indels for germline and cancer sequencing applications.□Bioinformatics. 2016;32(8):1220–1222.

31. Chen K, Wallis JW, McLellan MD, et al. BreakDancer: an algorithm for high-resolution mapping of genomic structural variation.□Nat Methods. 2009;6(9):677–681.

32. Ye K, Schulz MH, Long Q, et al. Pindel: a pattern growth approach to detect break points of large deletions and medium sized insertions from paired-end short reads.□Bioinformatics. 2009;25(21):2865–2871.

33. Mohiyuddin M, Mu JC, Li J, et al. MetaSV: an accurate and integrative structural-variant caller for next generation sequencing.□Bioinformatics. 2015;31(16):2741–2744.

34. Wala JA, Bandopadhayay P, Greenwald NF, et al. SvABA: genome-wide detection of structural variants and indels by local assembly.□Genome Res. 2018;28(4):581–591.

35. Wang S, Wang J, Xiao X, et al. GSDcreator: an efficient and comprehensive simulator for generating NGS data with population genetic information. In: 2019 IEEE International Conference on Bioinformatics and Biomedicine (BIBM). IEEE; 2019: 1868–1875.

36. Elliott M J, Howarth K, Main S, et al. Ultrasensitive detection and monitoring of circulating tumor DNA using structural variants in early-stage breast cancer[J]. Clinical Cancer Research, 2025, 31(8): 1520–1532.

37. Proietto M, Crippa M, Damiani C, et al. Tumor heterogeneity: preclinical models, emerging technologies, and future applications. Front Oncol. 2023; 13: 1164535.

38. Smolka M, Paulin LF, Grochowski CM, et al. Detection of mosaic and population-level structural variants with Sniffles2. Nat Biotechnol. 2024: 1–10.

39. Wang S, Zhu X, Wang X, et al. TMBstable: a variant caller controls performance variation across heterogeneous sequencing samples. Brief Bioinform. 2024; 25(3): bbae159.

40. Liu Z, Roberts R, Mercer TR, et al. Towards accurate and reliable resolution of structural variants for clinical diagnosis. Genome Biol. 2022; 23(1): 68.

41. Luijken K, Groenwold R H H, Van Calster B, et al. Impact of predictor measurement heterogeneity across settings on the performance of prediction models: A measurement error perspective[J]. Statistics in medicine, 2019, 38(18): 3444–3459.

42. Riley R D, Pate A, Dhiman P, et al. Clinical prediction models and the multiverse of madness[J]. BMC medicine, 2023, 21(1): 502.

