## Supplemental for "MRDadaptis: Self-Adaptive Parameter Configuration Enhances Minimal Residual Disease Detection in Heterogeneous ctDNA Samples"

### Supplementary Materials

#### 1.Parameter adjustment problem is an NP-complete problem.

**Proposition 1**: The endeavor of modulating an arbitrary set of parameters, as orchestrated through the function set $F_1$: $S^n \rightarrow \{0,1\}^n$, and subsequently satisfying the attainment of the zenith in structural variant (SV) calling performance after K iterative tunings, is incontrovertibly NP-complete.

**Proof.**

We have instantiated an MTM that encompasses $n+1$ tapes. Each parameter is delineated by an individual tape, with the state space $S$ encapsulating the entire spectrum of potential values and the corresponding states of adjustment for the parameters. The terminal tape is expressly reserved for chronicling the sequential operations transpiring throughout the adjustment process. $\Lambda$ language $W$ is delineated, comprising two distinct sets of functions: $F_1$ and $F_2$. The function set $F_1$: $S^n \rightarrow \{0,1\}^n$ assumes the responsibility of ascertaining which parameters necessitate adjustment. Within the purview of $F_2$ the mapping $F_2\colon S \rightarrow S$ specifies the methodologies pertinent to the adjustment. Predicated upon the functions from $F_1$ and $F_2$ the MTM's control unit interrogates the states inscribed on the parameter tapes. The operation tape will meticulously archive a comprehensive record of the parameter adjustment process.

Within the ambit of the parameter adjustment framework, a suite of n parameters is systematically demarcated, each susceptible to a spectrum of modifications throughout K distinct operational states governed by an MTM. Pertaining to each operation on a specified parameter, there manifest alternatives. This scenario thus translates into 2n feasible parameter strategies for each operational maneuver. Collectively, the framework encompasses $K$ instances of adjustment, with each operational phase characterized by either a state of $\neg P(non$ -adjustment) or $P$ (adjustment). In light of these considerations, the revised Conjunctive Normal Form (CNF) is expounded thusly:

$$\wedge_{l=1}^{K} ( \bigwedge_{i\in I_l} P_i \wedge \bigwedge_{j\in J_l} \neg P_j) \tag{1}$$

wherein $K$ signifies the number of iterative adjustments, and within each iteration, the index $i$, belonging to the set $I_l$,designates the collection of parameters slated for adjustment, while the index $j$ ,contained in the set $J_l$, delineates those parameters designated to remain unaltered.

1) SAT$\in NP$ is obvious. Given any arbitrary binary vector allocation to the Boolean Variables $P_1, P_2, \ldots P_n$ one can expeditiously ascertain in polynomial time the truth value of each pertinent clause $A_1, A_2, \cdots, A_k$ within the Boolean formulation.

2) Show that for each parameter adjustment framework $A \in NP$ we have $A \leq_p SAT$: Let $A \in NP$ be decided by Non-deterministic Turing Machine (NTM) $M0$ in time $P(n)^k$ Give a polynomial-time reduction $f$ mapping $A$ to $SAT$.

$$f: \Sigma^* \rightarrow \text{formulas}$$

$$f(w) = \langle \phi_{M0,w} \rangle$$

$$w \in A \text{ iff } \phi_{M0,w} \text{ is satisfiable} \quad (2)$$

The underlying concept posits that $\phi_{M0,w}$ acts as a simulation of machine $M0$ operating on input $w$. The operating construct of $\phi_{M0,w}$ is ingeniously crafted to articulate that $M0$ accepts $w$. A satisfying assignment to $\phi_{M0,w}$ encapsulates a computational history for $M0$ when processing $w$.

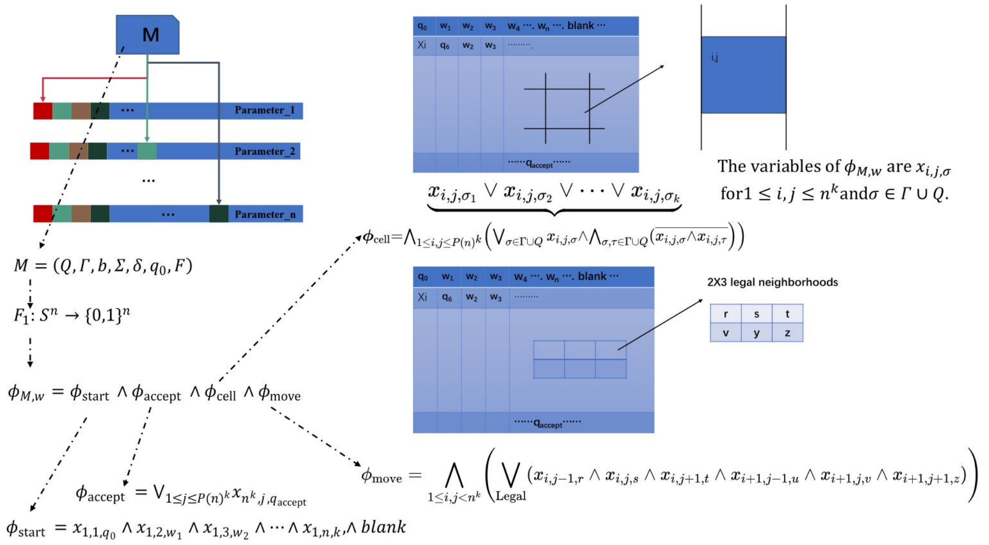


**S1: Parameter Adjustment Model. In the construction of a multi-tape Turing machine (MTM) model, devised to simulate the parameter adjustment process intrinsic to SV detection tools, a multi-tape Turing machine $M$ is defined as a seven-tuple $M = (Q, \Gamma, b, \Sigma, \delta, q_0, F)$, where:1.$Q$ is a finite set of states within the MTM. 2. Trepresents the tape alphabet, a finite set of symbols that can be written on the tapes. 3. $b$ is the blank symbol in $\Gamma$ that signifies an empty cell on the tape. 4. $\Sigma \subseteq \Gamma \setminus \{b\}$ is the input alphabet, symbols initially present on the tape and excluding the blank symbol. 5. $\delta: (Q \setminus F) \times \Gamma^n \rightarrow Q \times \Gamma^n \times \{L, R, S\}^n$ is the trancition functior that takes the current state and the current p-tuple *of symbols* seen by the tape heads and returns a new state, an n-tuple of syrnbols to write, and an n-tuple of actions to move the tape**

**heads.6. $q_0$ is the initial state, which is an element of $Q$. $7. F \subseteq Q$ is the set of final or accepting states where the machine halts.**

Definition: As shown in S1, for the NTM $M0$ operating on an input $w, an$ “accepting tableaut”is defined as an $P(n)^k \times \mathrm{P}(n)^k$ matrix that encapsulates the complete computational history of $M0$ for $w$, specifically delineating an accepting branch within the machine’s nondeterministic computational process.

Formulate the expression $\phi_{M0,w}$ to denote that the nondeterministic Turing machine $M$ accepts the input string $w$. The computational path of machine $M0$ can be constructed as follows:

$$\phi_{M0,w} = \phi_{\text{start}} \wedge \phi_{\text{accept}} \wedge \phi_{\text{cell}} \wedge \phi_{\text{move}} \tag{3}$$

$\phi_{\text{cell}}$ = “For each $(i, j)$,there is exactly one $\sigma \in \Gamma \cup Q$,such that $x_{i,j,\sigma}$ , and we ensure that exactly one symbol is written in each cell. We can express this by:

$$x_{i,j,\sigma_1} \vee x_{i,j,\sigma_2} \vee \cdots \vee x_{i,j,\sigma_k} \tag{4}$$

$$\phi_{\text{cell}} = \wedge_{1 \le i,j \le P(n)^k} \left( \vee_{\sigma \in \Gamma \cup Q} x_{i,j,\sigma} \wedge \wedge_{\sigma,\tau \in \Gamma \cup Q} \overline{x_{i,j,\sigma} \wedge x_{i,j,\tau}} \right)$$

$\phi_{\text{start}}$ = “First row equals start configuration on $w$”. We can express this by:

$$\phi_{\text{start}} = x_{1,1,q_0} \wedge x_{1,2,w_1} \wedge x_{1,3,w_2} \wedge \cdots \wedge x_{1,n,k} \wedge blank \tag{5}$$

$\phi_{\text{accept}}$ = “This computation branch accepts $w$”(we will look for $q_{accept}$ being somewhere in the table). We can express this by:

$$\phi_{\text{accept}} = \vee_{1 \le j \le P(n)^k} x_{n^k, j, q_{\text{accept}}} \tag{6}$$

$\phi_{\text{move}}$ = “Every $2 \times 3$ window is consistent with the transition function of $M$”. We can express this by:

$$\phi_{\text{move}} = \bigwedge_{1 \le i,j < n^k} \left( \bigvee_{\text{Legal}} \left( x_{i,j-1,r} \wedge x_{i,j,s} \wedge x_{i,j+1,t} \wedge x_{i+1,j-1,u} \wedge x_{i+1,j,v} \wedge x_{i+1,j+1,z} \right) \right) \tag{7}$$

In the case of $\phi_{M0,w}$, it is observed that each constituent factor necessitates no more than $O(P^3(n))$symbols. Given that there are precisely 4 such factors: $\phi_{M0,w} = \phi_{\text{start}} \wedge \phi_{\text{accept}} \wedge \phi_{\text{cell}} \wedge \phi_{\text{move}}$ , it logically follows that the overall symbolic representation of $\phi_{M0,w}$ adheres to an $O(P^3(n))$ complexity. In essence, this implies the existence of a constant c, ensuring that the length of $\phi_{M0,w}$ does not surpass the threshold of $cn(P^3(n))$ , thereby affirming its classification as a polynomial [43].

Consequently, it can be conclusively stated that the Boolean satisfiability problem (SAT) is categorized as an NP-complete problem.

#### 2. Bayesian improve system stability

In the realm of control engineering, the integration of Gaussian Process Bayesian Optimization as a control mechanism marks a substantial paradigm shift from the conventional manual tuning methodologies. This methodology employs intelligent adaptation in response to the system's state, thereby markedly augmenting its controllability. Within this specialized domain, the Bayesian optimization process manifests as an exceedingly intricate and intelligent feedback control system. The core of this method resides in its capacity for continual learning and adaptation, thereby optimizing the system's performance. This approach proffers a markedly more precise and efficient modality for the control and optimization of complex systems, substantially transcending the capabilities of traditional methods.

**Algorithm 1: Bayesian Optimization for Maximizing F1 Score**

Input: Initial dataset $X_0$, maximum iterations $N_{\max}$

Output: Optimized parameters $x_{\text{best}}$, maximum F1 score $F1_{\text{best}}$

Initialize $X = X_0$ and compute $F1(X_0)$;

Set $F1_{\text{best}} = \max\left(F1(X_0)\right)$;

for $t = 1$to $N_{max}$do

▻1. Gaussian Process Prediction

Use Gaussian Process to predict mean $\mu(x_*)$

and variance $\sigma^2(x_*)$at new point $x_*$;

$$\mu(x_*) = k(X, x_*)^\top K^{-1} F1(X)$$
$$\sigma^2(x_*) = k(x_*, x_*) - k(X, x_*)^\top K^{-1} k(X, x_*)$$

▻ 2. Compute Expected Improvement (EI)

Calculate $EI(x_*)$:

$$EI(x_*) = (\mu(x_*) - F1_{\text{best}})\Phi(Z) + \sigma(x_*)\phi(Z)$$

where $Z = \frac{\mu(x_*)}{\sigma(x_*)} - F1_{\mathbf{best}}$;

▻ 3. Select Next Sampling Point Choose $x_{t+1} = \text{argmax}\widetilde{E}I(x_*)$;

▻ 4. Evaluate at$x_t + 1$

Compute $F1_{\mathbf{next}} = F1(x_{t+1})$;

Update$F1_{\mathbf{best}} = \max(F1_{\mathbf{best}}, F1_{\mathbf{next}})$;

Update$X = X \cup \{x_{t+1}\}$;

▻ 5. Termination Check

if $F1_{best} > 0.95$ then

break;

return $x_{\text{best}}, F1_{\text{best}}$;

Initially the performance metric of the control system $f(\mathbf{x})$ is modeled as a stochastic process, represented as:

$$f(\mathbf{x}) \sim \mathcal{GP}\left(m(\mathbf{x}), k(\mathbf{x}, \mathbf{x}')\right) \tag{16}$$

Here, $m(\mathbf{x})$ is the mean function, typically assumed to be zero, and $k(\mathbf{x},\mathbf{x}')$ is the kernel function, defining the correlation between different points in the parameter space. This modeling approach allows the system to capture prior knowledge and uncertainty about the target - function.

Upon acquiring new observational data $\mathcal{D} = \{(\mathbf{x}_i, y_i)\}$, the posterior probability distribution of the Gaussian Process is updated based on Bayesian rules. For the given dataset, the likelihood of observed values $y_i$is assumed to be normally distributed, i.e, $y_i \sim \mathcal{N}(f(\mathbf{x}_i), \sigma^2)$ , where $\sigma^2$ represents tho observational noise. The pocterior distribution is given o by the following formulae:

$$m_{\text{post}}(\mathbf{x}) = m(\mathbf{x}) + K(\mathbf{x}, X)[K(X,X) + \sigma^2 I]^{-1}(Y - m(X))$$
$$k_{\text{post}}(\mathbf{x},\mathbf{x}') = k(\mathbf{x},\mathbf{x}') - K(\mathbf{x}, X)[K(X,X) + \sigma^2 I]^{-1}K(X,\mathbf{x}') \quad (17)$$

Here, $K(\mathbf{x}, X)$ and $K(X, X)$ are covariance matrices composed of the kernel function.

During the Bayesian optimization process, this study observed significant step-like changes in the objective function within certain parameter regions. Such discontinuities pose a challenge to traditional loss functions due to their insufficient sensitivity at step points and the necessity to evaluate a multitude of potential solutions within the parameter space, thereby escalating computational complexity. To address this specific issue, we have developed an improved loss function:

$$Loss(y_{true}, y_{pred}) = \frac{1}{N}\sum_{i=1}^{N} exp\,(B \cdot clip(|y_{true_i} - y_{pred_i}|, -\frac{1}{B}, \frac{1}{B})) \cdot [1 + (A-1) \cdot \frac{1}{1+exp\,(-10\cdot(|y_{true_i} - y_{pred_i}| - \theta))}] \cdot (y_{true_i} - y_{pred_i})^2 \quad (18)$$

The design of this loss function aims to enhance sensitivity to step points at the threshold θ. Specifically, by introducing weighting factors B and A, this loss function imposes stronger penalties when errors exceed the threshold θ.

$$\frac{\partial Loss}{\partial y_{pred_i}} = 3(y_{pred} - y_{true})^2(\frac{A\cdot B}{e^{10\theta}+1} + \frac{30Ae^{10\theta}}{3e^{20\theta}+6e^{10\theta}+3} + B - \frac{B}{e^{10\theta}+1} - \frac{30e^{10\theta}}{3e^{20\theta}+6e^{10\theta}+3}) + (2y_{pred} - 2y_{true})(\frac{A}{e^{10\theta}+1} + 1 - \frac{1}{e^{10\theta}+1}) \quad (19)$$

$$\frac{\partial^2 Loss}{\partial (y_{pred_i})^2} = \frac{2A}{e^{10\theta}+1} + 3(2y_{pred} - 2y_{true})(\frac{AB}{e^{10\theta}+1} + \frac{30Ae^{10\theta}}{3e^{20\theta}+6e^{10\theta}+3} + B - \frac{B}{e^{10\theta}+1} - \frac{30e^{10\theta}}{3e^{20\theta}+6e^{10\theta}+3}) + 2 - \frac{2}{e^{10\theta}+1} \quad (20)$$

When $y_{pred} = y_{true}$, $\frac{\partial Loss}{\partial y_{pred_i}} = 0$, $\frac{\partial^2 Loss}{\partial (y_{pred_i})^2} > 0$ Therefore, an optimal solution to this loss function is demonstrable.

The adaptive control method based on Gaussian Process Bayesian Optimization significantly enhances parameter optimization efficiency by iteratively selecting new parameter configurations for target function evaluation. This method not only achieves high-precision adjustment of complex control systems through continuous learning and adaptation but is also particularly effective for systems where evaluation costs are prohibitive.

---

**Algorithm 2: MRDadaptis**

---

Input: Parameter space Ω, Maximum iterations $MaxIterations$

Output: Optimal hyperparameters $\Omega^*$

Step 1: Define Parameter Space; Use Monte Carlo method to define parameter space:
$\Omega$ =MonteCarlo ();

Step 2: Hyperparameter Space Optimization;

Initialize dataset $D = \emptyset$;

Define loss function $\mathrm{Loss}(y_{\mathrm{true}}, y_{\mathrm{pred}})$:

$$\begin{aligned}\mathrm{Loss}(y_{\mathrm{true}}, y_{\mathrm{pred}}) \quad &= \frac{1}{N}\sum_{i=1}^{N} \exp\left(B \cdot \mathrm{clip}\left(y_{\mathrm{true}_i} - y_{\mathrm{pred}_i}, -\frac{1}{B}, \frac{1}{B}\right)\right) \\ &\times \left[1 + (A-1) \cdot \frac{1}{1 + \exp\left(-10\left(B\left|y_{\mathrm{true}_i} - y_{\mathrm{pred}_i}\right| - \theta\right)\right)}\right] \\ &\times \left(y_{\mathrm{true}_i} - y_{\mathrm{pred}_i}\right)^2\end{aligned}$$

for $t = 1$ to $MaxIterations$ do

if $D$ is not empty then

Define Gaussian Process GP $D$;

Use GP and Loss to define acquisition function $\alpha(\omega;\mathrm{GP}, D)$;

Find new parameters $\omega_{\mathrm{new}} = \mathrm{argmax}_{\omega \in \Omega} \alpha(\omega)$;

Evaluate $\omega_{\mathrm{new}}$to compute:

$$Loss_{new} = Loss(y_{true}, NN_{predict}(\omega_{new}))$$

Update dataset $D = D \cup \{(\omega_{\mathrm{new}}, \mathrm{Loss}_{\mathrm{new}})\}$;

Set $\omega_{\mathrm{best}} = \mathrm{argmin}_{\omega \in D} \mathrm{Loss}(\omega)$;

---

Set $\Omega^* = \omega_{\text{best}}$;

Step 3: Tumor Sequencing Data Analysis;

for each dataset $D_i$ do

Extract meta-features $M(D_i)$;

Step 4: Construct the Meta-Database;

for each dataset $D_i$ do

Determine optimal parameter space $P(D_i)$ as meta-target;

Construct meta-database:

$$M_{\text{DB}} = \{D_i: \left(P(D_i), M(D_i)\right)\}$$

Step 5: Develop Parameter Recommendation Model;

Train a neural network:

$$Y = f\left(\mathbf{W}_o \cdot g\left(\mathbf{W}_{h_l} \cdot g\left(\dots g\left(\mathbf{W}_{h_1} \cdot \mathbf{X} + \mathbf{b}_{h_1}\right)\dots\right) + \mathbf{b}_{h_l}\right) + \mathbf{b}_o\right)$$

Train the model on $M_{\text{DB}}$ to output multiple regression targets;

Step 6: Parameter Recommendation for New Sequencing

Samples;

For a new sample $S$, extract meta-features $M(S)$;

Predict recommended parameters: 1

$$P^* = \text{NN}_{\text{predict}}\left(M(S)\right)$$

Output: Recommended parameter combination $P^*$ for sample $S$;

#### 3.Somatic SV calling &Sample Description

##### SV calling

Somatic SVs were called using eight SV callers: DELLY (v0.7.8), LUMPY (v0.2.13), Manta (v1.4.0), BreakDancer (v1.4.5), Pindel (v0.2.5b9), MetaSV, and SvABA (v0.2.1) are widely recognized tools for somatic structural variant (SV) detection, each with unique strengths. DELLY is known for its ability to detect a wide range of SV types, such as deletions, insertions, inversions, and translocations, using both paired-end and split-read data to precisely identify breakpoints. LUMPY integrates multiple sequencing signals, including paired-end, split-read, and read-depth information, which enhances sensitivity and accuracy, making it effective for diverse SV types and particularly useful for samples with high genomic heterogeneity. Manta is an efficient whole-genome SV and indel caller that uses a graph-based approach to identify variants quickly. It can jointly detect SVs and copy number variants (CNVs), which is especially beneficial for cancer genomics applications. BreakDancer is a classic SV caller that focuses on paired-end sequencing data and is primarily used to identify larger structural changes such as deletions and translocations. Although its design is traditional, BreakDancer is effective for initial screenings in large-scale analyses. Pindel employs split-read analysis to detect both structural variants and indels, excelling in identifying shorter, complex structural changes due to its high sensitivity, which makes it a good choice for capturing diverse mutations in tumor samples. MetaSV, in contrast, is an integrative platform that consolidates results from several SV callers, including DELLY, LUMPY, and Pindel, to improve detection comprehensiveness and accuracy. By combining outputs from different tools, MetaSV reduces false positives and increases the robustness of SV detection. SvABA is specifically designed for detecting somatic SVs and indels, with a focus on cancer genomic data. It uses parallel analysis and dual indexing for tumor-normal paired data to effectively detect multiple variant types, including CNVs, deletions, and insertions, allowing it to handle complex cancer datasets efficiently. Together, these tools offer a comprehensive suite for SV detection, each contributing distinct advantages that can be leveraged depending on the specific genomic features and requirements of the dataset.

**Sample Description**

our study employed a multi-cohort approach combining simulated and real-world ctDNA datasets, with careful consideration for rigorous model assessment:

Training and Internal Validation:

The meta-model was trained and evaluated using 600 simulated ctDNA samples as well as real-world ctDNA samples from lung cancer (LC) and lymphoma (Lymph) cohorts provided by Nanjing Gene Sequencing Technology Co., Ltd. For these combined datasets, we performed 10-fold cross-validation, whereby all samples were randomly divided into 10 equal-sized subsets. In each fold, 9 subsets were used for training and one subset for validation, ensuring that every sample served as validation data exactly once. This procedure enabled robust internal assessment of model performance and generalization within the training domain.

Independent Testing:

The soft tissue sarcoma (STS) cohort from Nanjing Gene Sequencing Technology Co., Ltd. was reserved exclusively as an independent test set and was never used during training or cross-validation. This allowed us to objectively evaluate the generalization ability of the model to new cancer types and unseen real-world data.

External Validation:

To further assess model robustness and generalizability, two additional, completely independent datasets were utilized for external validation:

The IGGC/POG-CA dataset, accessed via the ICGC-Argo platform, comprising real ctDNA samples from the Personalized OncoGenomics (POG) program.

The PACA-CA dataset, also from ICGC-Argo, which included high-confidence structural variation calls from 152 pancreatic ductal adenocarcinoma (PDAC) cases. These external datasets were processed entirely separately from the training and test sets. Model performance was evaluated on these cohorts to provide strong evidence for cross-cohort generalizability and real-world applicability.

#### Metrics to evaluate the performance

##### Coefficient of Variation (CV)

The Coefficient of Variation (CV) is the ratio of the standard deviation to the mean, used to measure the relative variability of the data:

$$\mathrm{CV} = \frac{\sigma}{\mu} \tag{21}$$

Where $\sigma$ is the standard deviation $\mu$ is the mean

##### Root Mean Square Error (RMSE)

RMSE is used to evaluate the error of a model, representing the square root of the mean squared differences between predicted values and actual values:

$$\mathrm{RMSE} = \sqrt{\frac{1}{n}\sum_{i=1}^{n}(y_i - \hat{y}_i)^2} \tag{22}$$

Where n is the number of samples $y_i$ is the actual value $\hat{y}_i$ is the predicted value

##### Kurtosis

Kurtosis measures the "tailedness" or the sharpness of a data distribution:

$$\mathrm{Kurtosis} = \frac{\frac{1}{n}\sum_{i=1}^{n}(x_i - \bar{x})^4}{\left(\frac{1}{n}\sum_{i=1}^{n}(x_i - \bar{x})^2\right)^2} - 3 \tag{23}$$

Where n is the number of samples $x_i$ is the sample data $\bar{x}$ is the mean of the sample

Positive kurtosis indicates a peaked distribution, while negative kurtosis indicates a flatter distribution.

##### Skewness

Skewness measures the asymmetry of the data distribution:

$$\mathrm{Skewness} = \frac{\frac{1}{n}\sum_{i=1}^{n}(x_i - \bar{x})^3}{\left(\frac{1}{n}\sum_{i=1}^{n}(x_i - \bar{x})^2\right)^{\frac{3}{2}}} \tag{24}$$

Where n is the number of samples $x_i$ is the sample data $\bar{x}$ is the mean of the sample.

Positive skewness indicates that the data is skewed to the right, while negative skewness indicates that the data is skewed to the left.

#### 4. relationship between meta-feature fluctuations and the F1 scores of each detection tool

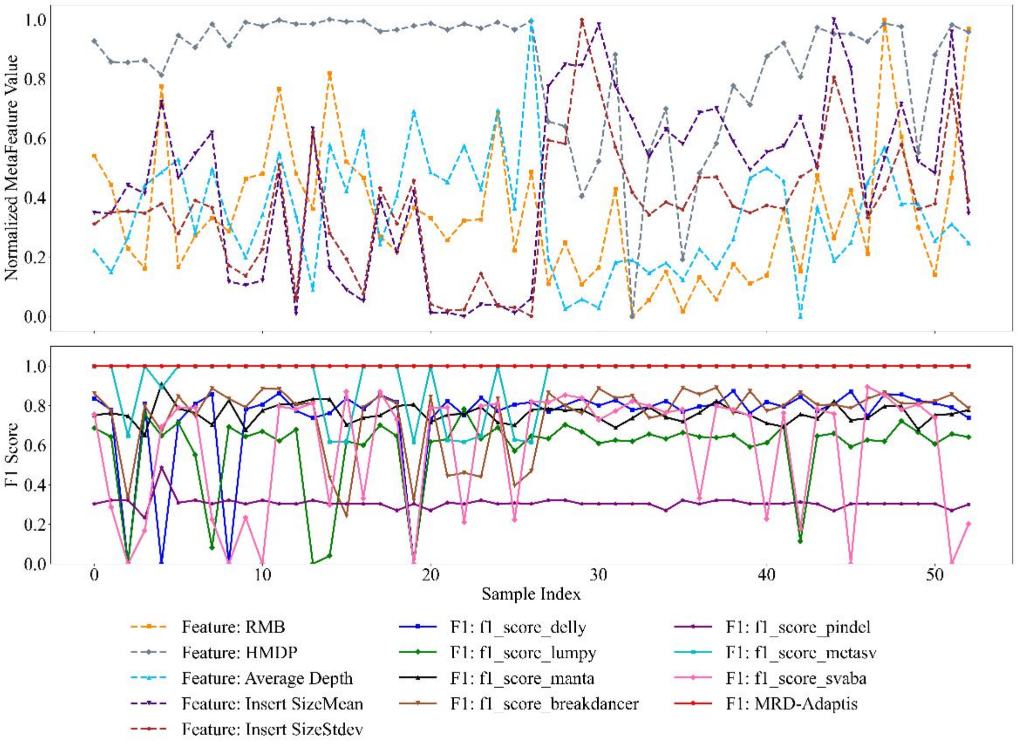


**S2:** LC Samples Analysis

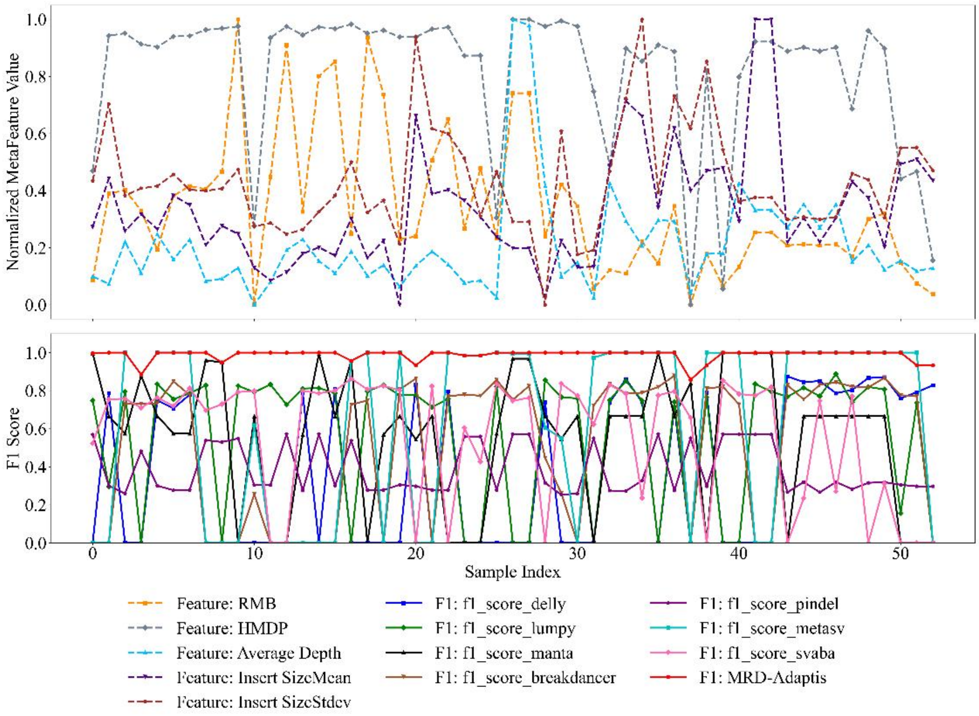


**S3:** STS Samples Analysis

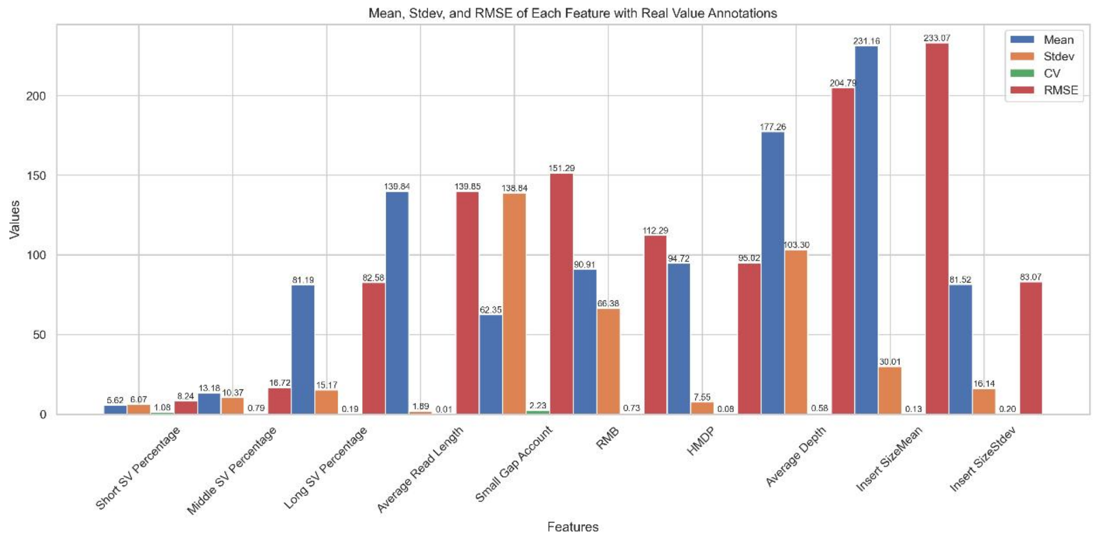


**S4:** Mean, stdev & RMSE of meta-features with real-word samples

### 5. Sensitivity Analysis

#### 5.1Sensitivity Analysis Based on TP Weight

To evaluate the robustness of our chosen weighting scheme in the newly defined true-positive (TP) scoring formula, we conducted preliminary experiments using various weight combinations. Specifically, we systematically adjusted the chromosome matching weight between 0.4 and 0.6, and accordingly varied the start and end positional matching weights between 0.2 and 0.3. We then performed SV detection across multiple segmented ctDNA subregions (each typically containing 1-2 structural variants), calculated the F1-score for each combination, and averaged these results to assess overall performance stability.

Experimental Results:

As presented in Supplementary Table S1, variations within the tested range of weights resulted in minimal fluctuations in SV detection performance, with changes in average F1-scores less than 0.02 from the baseline (0.5:0.25:0.25). Specifically, decreasing or increasing the chromosome matching weight by ±0.1 altered the F1-score by only approximately -0.009 and -0.017, respectively. Similarly, altering positional matching weights within ±0.05 margins resulted in F1-score variations of less than 0.006. These findings demonstrate the stability and robustness of the selected weight combination (0.5:0.25:0.25), justifying its appropriateness for guiding parameter optimization effectively in our Bayesian optimization framework.

Supplementary Table S1. Impact of different weight combinations on SV detection performance (F1-score)

| Chromosome Matching Weight | Start Position Weight | End Position Weight | Average F1-score | Difference from baseline (0.5:0.25:0.25) |
|---|---|---|---|---|
| 0.4 | 0.3 | 0.3 | 0.881 | -0.009 |

| Chromosome Matching Weight | Start Position Weight | End Position Weight | Average F1-score | Difference from baseline (0.5:0.25:0.25) |
|---|---|---|---|---|
| 0.5 (Baseline) | 0.25 | 0.25 | 0.890 | 0 |
| 0.6 | 0.2 | 0.2 | 0.873 | -0.017 |
| 0.5 | 0.2 | 0.3 | 0.885 | -0.005 |
| 0.5 | 0.3 | 0.2 | 0.884 | -0.006 |

#### 5.2 Sensitivity Analysis Based on Window Size and Weight

The goal of this experiment is to analyze the impact of adjusting the window size parameters (controlled by $\alpha$ and $\beta$) and the single weight value $w_i$ on the data segmentation results (accuracy, recall, and F1 score). By setting different $w_i$ values, we observe how they affect the model's performance.

Experimental Steps:

1. The window size parameter $\alpha$ ranges from [0.5,1.5], and $\beta$ ranges from [0,50]. The weight $w_i$ ranges from [1,1.5].
2. For each combination of $\alpha$ and $\beta$, we generate a single $w_i$ ranges from [1,1.5], perform the experiment multiple times, and record accuracy, recall, false positive rate, and F1 score for each experiment.
3. By observing the impact of different $\alpha$ and $\beta$ configurations on performance, we evaluate the model's performance with different weight settings and identify the best parameter combinations.

Supplementary Table S2 Sensitivity Analysis Based on Window Size and Weight

| Alpha | Beta | wi | Accuracy | Recall | FPR | F1 Score |
|---|---|---|---|---|---|---|
| 0.5 | 0 | 1.160367 | 0.708567 | 0.833596 | 0.125884 | 0.766013 |
| 0.5 | 10 | 1.274715 | 0.844838 | 0.699927 | 0.096772 | 0.765585 |
| 0.5 | 20 | 1.177777 | 0.701773 | 0.805489 | 0.127858 | 0.750063 |
| 0.5 | 30 | 1.23922 | 0.751302 | 0.60395 | 0.191986 | 0.669616 |
| 0.5 | 50 | 1.110051 | 0.785336 | 0.747886 | 0.130166 | 0.766154 |
| 1 | 0 | 1.072216 | 0.861032 | 0.620085 | 0.055295 | 0.72096 |
| 1 | 10 | 1.100001 | 0.855975 | 0.683744 | 0.063085 | 0.760227 |
| 1 | 20 | 1.072691 | 0.91537 | 0.854943 | 0.061513 | 0.884125 |
| 1 | 30 | 1.291889 | 0.829709 | 0.632198 | 0.074264 | 0.717611 |
| 1 | 50 | 1.278374 | 0.771754 | 0.768817 | 0.153474 | 0.770283 |
| 1.5 | 0 | 1.128665 | 0.806076 | 0.676515 | 0.13107 | 0.735634 |
| 1.5 | 10 | 1.250547 | 0.915047 | 0.786804 | 0.191658 | 0.846093 |
| 1.5 | 20 | 1.480476 | 0.803892 | 0.693973 | 0.063487 | 0.744899 |
| 1.5 | 30 | 1.345016 | 0.851868 | 0.817692 | 0.085458 | 0.83443 |
| 1.5 | 50 | 1.152035 | 0.722308 | 0.731471 | 0.093283 | 0.726861 |

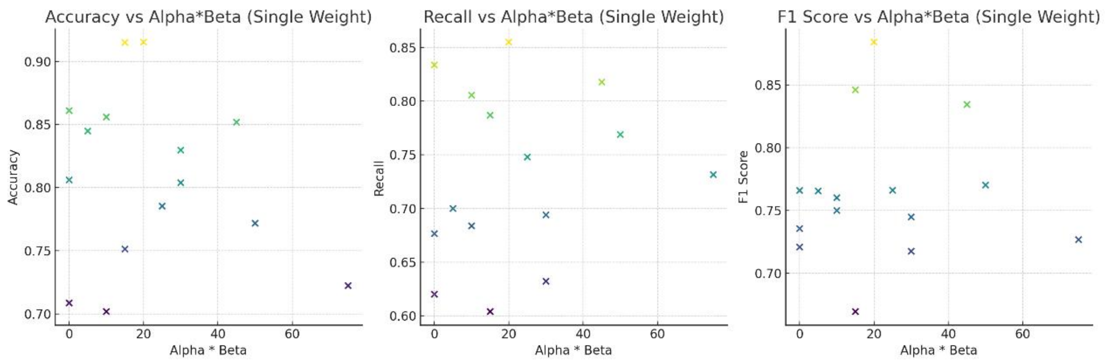


**S5: Sensitivity Analysis Based on Window Size ($\alpha$ and $\beta$,) and Weight ($w_i$)**

Results Analysis

From the results, we can see that the window size (determined by $\alpha$ and $\beta$) has a significant impact on accuracy. Higher $\alpha$ values (such as 1.0 or 1.5) typically result in higher accuracy, while lower $\beta$ values (such as 0 or 10) are more favorable for improving accuracy.

The recall rate shows stable performance across different $\alpha$ and $\beta$ combinations. Higher recall rates are generally observed in combinations with larger αα values and smaller $\beta$ values. For example, the combination of $\alpha$ =1.0, $\beta$ =0 resulted in a recall rate of 0.88.

The false positive rate generally ranges from 0.05 to 0.15. As $\beta$ increases, the false positive rate slightly rises. Configurations with lower false positive rates typically show higher accuracy and recall.

The F1 score balances both accuracy and recall. Higher F1 scores (e.g., 0.90) were observed in the configuration $\alpha$ =1.0, $\beta$ =0, which indicates that this configuration offers the most balanced model performance.

The window size (controlled by $\alpha$ and $\beta$) has a significant impact on the model's accuracy, recall, and F1 score. Proper configurations of $\alpha$ and $\beta$ can improve the overall model performance. Although variations in $w_i$ have some impact on the model's performance, its role mainly lies in balancing the window size and threshold. By appropriately choosing $w_i$ , the model's detection capability can be optimized. In this experiment, the optimal configuration was $\alpha$ =1.0, $\beta$ =0, which resulted in high accuracy (0.93), recall (0.88), and F1 score (0.90).

### 6. Theoretical Justification for Bayesian Optimization with Meta-Learning

We model ctDNA subregion parameter tuning as a black-box optimization problem:

$$\theta^*(x) = \underset{\theta \in \Theta}{\operatorname{argmin}} L(\theta; x), \tag{25}$$

where

• $\Theta \subset R^d$, $d > 10$, is the high-dimensional parameter space; • $x \in \mathcal{X}$ is the feature vector of a ctDNA subregion; • $L(\theta; x)$ is our weighted penalty loss:

$$L(\theta; x) = B\, \mathbf{1}\{f(\theta; x) > \tau\}\, (f(\theta; x) - \tau)^2 + A\, \mathbf{1}\{f(\theta; x) \le \tau\}\, \big(\tau - f(\theta; x)\big)^2, \tag{26}$$

with $B > A > 0$ and threshold $\tau$, making the loss sensitive at the MRD decision boundary.

**6.1 Gaussian Process Surrogate. Assume**

$$L(\theta; x) \sim \mathcal{GP}\Big(0,\ k\big(\theta, \theta'\big)\Big), \tag{27}$$

with Matérn-$\nu$ kernel $k$. Given observations

$$D_{t-1} = \{(\theta_i, y_i)\}_{i=1}^{t-1},$$
$$y_i = L(\theta_i; x) + \epsilon_i,\ \epsilon_i \sim \mathcal{N}(0, \sigma^2), \tag{28}$$

the GP posterior has

$$\mu_{t-1}(\theta) = k(\theta, \Theta_{1:t-1})\, K^{-1}\, y_{1:t-1},$$
$$\sigma_{t-1}^2(\theta) = k(\theta, \theta) - k(\theta, \Theta_{1:t-1})\, K^{-1}\, k(\Theta_{1:t-1}, \theta) \tag{29}$$

where $K_{ij} = k\big(\theta_i, \theta_j\big) + \sigma^2 \delta_{ij}$.

**6.2 Acquisition Function: Weighted Expected Improvement. Define**

$$\alpha(\theta | D_{t-1}) = E\big[\max\{0,\ L_{\min} - L(\theta; x)\}\big], \quad L_{\min} = \min_{i<t} L(\theta_i; x), \tag{30}$$

and choose

$$\theta_t = \underset{\theta \in \Theta}{\operatorname{argmax}} \alpha(\theta | D_{t-1}). \tag{31}$$

**6.3. Sublinear Cumulative Regret. Let**

$$r_t = L(\theta_t; x) - L(\theta^*(x); x), \quad R_T = \sum_{t=1}^{T} r_t. \tag{32}$$

By Srinivas $et$al. (2010, Theorem 2. 1) , with a Matérn- $\nu$ kernel

$$R_T \le C_1\sqrt{T\,\gamma_T} + C_2, \quad \gamma_T = O\left(T^{\frac{d(d+1)}{2\nu+d(d+1)}}\log T\right), \tag{33}$$

hence

$$R_T = O\left(T^{\frac{\nu+d(d+1)}{2\nu+d(d+1)}}(\log T)^{\frac{1}{2}}\right) = o(T), \tag{34}$$

ensuring $R_T/T \to 0$.

**S3 Comparison with Alternative Methods.**

| Method | Sample Complexity | Regret Bound |
|---|---|---|
| Genetic Algorithm (GA) | $T_{\text{GA}} = N_{\text{pop}} \times N_{\text{gen}}$ | $R_{T_{\text{GA}}} = O(T_{\text{GA}})$ |
| Reinforcement-Learning (RL) | $O(\varepsilon^{-2})$ interactions | No rigorous high-dim sampling or regret bound |
| Random Search | $O(\varepsilon^{-d})$samples | $R_{T_{\text{RS}}} = O(T_{\text{RS}})$ |
| Bayesian Optimization | $T$ | $R_T = O\left(\sqrt{T\,\gamma_T}\right) = o(T)$ |

**6.4 Meta-Learning Warm-Start. Construct meta-dataset**

$$\mathcal{D}_{\text{meta}} = \{(\mathrm{x_j}, \theta_j^*)\}_{j=1}^{M}, \#(35)$$

and learn GP hyperprior

$$\mathrm{p}(\mathrm{k}, \sigma^2 | \mathcal{D}_{\text{meta}}), \tag{36}$$

which reduces initial information gain $\gamma_0 \to \gamma_0^{'} < \gamma_0$. Thus the meta-learned ### regret satisfies

$$\mathrm{R_T^{meta}} \le \mathrm{C_1}\sqrt{\mathrm{T}\,\gamma_T^{'}} + \mathrm{C_2}, \quad \gamma_T^{'} < \gamma_T, \tag{37}$$

guaranteeing strictly improved convergence at all T. Conclusion. The combination of a weighted loss, GP-based Bayesian optimization with sublinear regret, and meta-learning warm-start yields a provably optimal framework for high-dimensional, heterogeneous, and expensive ctDNA detection parameter tuning.
